# Lytic Reactivation of the Kaposi’s sarcoma-associated herpesvirus (KSHV) is Accompanied by Major Nucleolar Alterations

**DOI:** 10.1101/2020.05.15.097808

**Authors:** Nofar Atari, K. Shanmugha Rajan, Vaibhav Chikne, Smadar Cohen-Chalamish, Odelia Orbaum, Avi Jacob, Inna Kalt, Shulamit Michaeli, Ronit Sarid

## Abstract

The nucleolus is a sub-nuclear compartment whose primary function is the biogenesis of ribosomal subunits. Certain viral infections affect the morphology and composition of the nucleolar compartment and influence ribosomal RNA (rRNA) transcription and maturation. However, no description of nucleolar morphology and function during infection with Kaposi’s sarcoma-associated herpesvirus (KSHV) is available to date. Using immunofluorescence microscopy, we documented extensive destruction of the nuclear and nucleolar architecture during lytic reactivation of KSHV. This was manifested by redistribution of key nucleolar proteins, including the rRNA transcription factor, UBF, the essential pre-rRNA processing factor Fibrillarin, and the nucleolar multifunctional phosphoproteins Nucleophosmin (NPM1) and Nucleolin. Distinct delocalization patterns were evident; certain nucleolar proteins remained together whereas others dissociated, implying that nucleolar proteins undergo nonrandom programmed dispersion. Of note, neither Fibrillarin nor UBF colocalized with promyelocytic leukemia (PML) nuclear bodies or with the viral protein LANA-1, and their redistribution was not dependent on viral DNA replication or late viral gene expression. No significant changes in pre-rRNA levels and no accumulation of pre-rRNA intermediates were found by RT-qPCR and Northern blot analysis, respectively. Furthermore, fluorescent *in situ* hybridization (FISH), combined with immunofluorescence, revealed an overlap between Fibrillarin and internal transcribed spacer 1 (ITS1), which represents the primary product of the pre-rRNA, suggesting that the processing of rRNA proceeds during lytic reactivation. Finally, small changes in the levels of pseudouridylation were documented across the rRNA. Taken together, our results suggest that rRNA transcription and processing persist during lytic reactivation of KSHV, yet they may become uncoupled. Whether the observed nucleolar alterations favor productive infection or signify cellular anti-viral responses remains to be determined.

**Author Summary:** We describe the extensive destruction of the nuclear and nucleolar architecture during lytic reactivation of KSHV. Distinct delocalization patterns are illustrated: certain nucleolar proteins remained associated with each other whereas others dissociated, implying that nucleolar proteins undergo nonrandom programmed dispersion. Of note, no significant changes in pre-rRNA levels and no accumulation of pre-rRNA intermediates were found, suggesting that pre-RNA transcription and processing continue and could be uncoupled during lytic reactivation. Small changes in the levels of pseudouridylation were documented across the rRNA. Previous studies showed that the different forms of KSHV infection are controlled through cellular and viral functions, which reprogram host epigenetic, transcriptomic, post-transcriptomic and proteomic landscapes. The ability of KSHV to affect the nucleolus and rRNA modifications constitutes a novel interaction network between viral and cellular components. The study of rRNA modifications is still in its infancy; however, the notion of altering cell fate by regulating rRNA modifications has recently begun to emerge, and its significance in viral infection is intriguing.

## Introduction

KSHV, also known as human herpesvirus 8 (HHV-8), is a cancer-related gamma-2 herpesvirus which is etiologically implicated in all types of Kaposi’s sarcoma (KS). In addition, KSHV is the causative agent of other disorders, including primary effusion lymphoma (PEL), multicentric Castleman’s disease, and KSHV-inflammatory cytokine syndrome [1-6]. Like all herpesviruses, KSHV undergoes either lytic (productive) or latent infection. During latent infection, the virus becomes largely transcriptionally quiescent and no virions are produced. In contrast, the productive course involves extensive transcription, replication, assembly and packaging of viral DNA, and ends with the release of new virions. The productive infection takes place in the host cell nucleus and is accompanied by drastic reorganization of the nuclear architecture involving conformational changes of the nuclear lamina and positioning of condensed chromatin at the nuclear periphery. At the same time, globular viral replication compartments (RCs), devoid of chromatin, are formed and then merge into a kidney-shaped zone characterized by high levels of viral DNA synthesis and high concentrations of viral and host proteins. Viral transcriptional factories, which recruit a significant fraction of RNA polymerase II, also accumulate in the nucleus, mostly in foci in and around RCs [7-9]. These events have been comprehensively studied for certain herpesviruses [10-13], yet the characteristics of nuclear alterations, particularly those in the nucleolus, during KSHV infection have only begun to be elucidated.

The nucleolus is a non-membrane bound sub-nuclear structure whose primary function is the biogenesis of ribosomal subunits, accomplished by the transcription of pre-ribosomal RNA (pre-rRNA), rRNA processing and assembly with ribosomal proteins. These functions are reflected by three distinct nucleolar sub-structures known as the fibrillar center (FC), the dense fibrillar component (DFC), and the granular component (GS). The nucleolus is surrounded by heterochromatin, forming perinucleolar chromatin [14-18]. The FC is rich in upstream binding factor (UBF), which plays a role in the recruitment of RNA polymerase I to rDNA promoters, and in chromatin structure modulation. Transcription of pre-rRNA takes place either in the FC or at the boundary between the FC and the DFC. Post-transcriptional processing and modifications of pre-rRNA occur in the DFC region, which contains processing factors such as the small nucleolar-associated enzyme 2’-O-methyltransferase Fibrillarin, and the pseudouridine synthase Dyskerin (DKC1). Late rRNA processing, along with assembly and transport of ribosomal subunits, occurs in the GC region, which comprises the remainder of the nucleolus and is enriched in processing and assembly factors such as Nucleophosmin 1 (NPM1/B23) and Nucleolin, as well as ribosomal proteins. Of note, the nucleolus is also involved in other cellular processes that may not be directly associated with the biogenesis of ribosomal subunits, including cell cycle control, stress responses, senescence, telomerase activity, protein degradation and sequestration, chromosomal domain organization, and innate immune responses [19, 20]. Hence, the nucleolus has a dynamic composition of proteins and RNA molecules with diverse functions.

Viral infections may cause nucleolar alterations, manifested by the redistribution of host nucleolar molecules to different cellular sites, the occurrence of distinct RNA or protein modifications, and the targeting of cellular proteins to the nucleoli. In addition, viral products may traffic to and from the nucleolus, and thereby alter the cell cycle, mRNA transport and innate host cell immunity. A large proportion of these activities are not restricted to any particular type of virus [21-26]. For example, replication and transcription of Borna disease virus take place in the nucleolus [27], assembly of AAV occurs in the nucleolus [28], REV protein of HIV is localized to the nucleolus [29, 30], and modifications of the basal rRNA transcription factors, Selectivity Factor-1 (SL-1) and UBF, inhibit the activity of RNA polymerase I during poliovirus infection [31, 32]. In addition, NSs protein encoded by Schmallenberg virus of the bunyaviridae family, is targeted to the nucleolus and triggers nucleolar disruption [33], and redistribution of UBF, Nucleolin and NPM1 during adenovirus infection promotes nuclear reorganization, viral DNA replication, and virion assembly [23, 34-39]. An almost complete destruction of the nucleolar structure, involving dispersion of nucleolar proteins, including UBF, Fibrillarin, Nucleolin and NPM1 to distinct cellular sites, has been described following infection with herpes simplex virus type 1 (HSV-1) [40, 41]. Redistribution of the nucleolar proteins following HSV-1 is partially dependent on the conserved late viral protein, UL24 [42-44]. Yet, it has been suggested that rRNA transcription and synthesis of ribosomal proteins, along with assembly of ribosomes, continue at a rate only slightly below that in uninfected cells [45, 46]. Nevertheless, an altered rRNA maturation pathway has been described in HSV-1-infected cells [47]. Of note, nucleolar proteins were found in proteomes that associate with viral genomes during infection with Adenovirus and HSV-1 [48].

Three cellular nucleolar proteins, including Angiogenin, NPM1 and Nucleolin, were shown to regulate KSHV infection, and three viral lytic gene products, ORF57, vBcl-2 and ORF20, localize to the nucleolus. Specifically, infection of endothelial cells with KSHV induces the expression and secretion of the angiogenesis factor Angiogenin, which in-turn translocates into the nucleus and to the nucleolus of subconfluent cells, stimulates proliferation and 45S rRNA gene transcription, and inhibits apoptosis [49]. vCyclin-Cdk6-mediated phosphorylation of NPM activates its association with LANA-1 to facilitate the interaction of LANA-1 with HDAC1 and core histones to sustain latency, while NPM knock-down leads to viral reactivation [50]. Binding of Nucleolin to IL-6 mRNA 3’ UTR was shown to promote escape from virus-induced degradation of host mRNAs [51]. ORF57 has major roles in transcriptional activation, RNA stability, RNA nuclear export and translational enhancement [52, 53]. vBcl-2, is a homolog of the Bcl-2 family of apoptosis and autophagy regulators [54-56], and is targeted to the nucleolus via interaction with the nucleolar protein PICT-1 [57]. ORF20 is a member of the UL24 protein family, which is conserved in all three subfamilies of the Herpesviridae. ORF20 co-purifies with 40S and 60S ribosomal subunits, and is associated with polysomes [58], however it is not yet known whether the association of ORF20 with ribosomal proteins is established in the nucleolus during ribosomal biogenesis or at later times.

In the present study, we describe major alterations in the nucleolar organization during lytic reactivation of KSHV-infected cells, involving redistribution of key nucleolar proteins which are known to be involved in the biogenesis of the ribosomal subunits. Yet, we provide evidence that pre-rRNA transcription proceeds and rRNA processing continues. Finally, we demonstrate that pseudouridylation, one of the predominant rRNA modifications, persists but with changes in the levels of this modification, likely affecting ribosome function during virus propagation.

## Results

### Lytic reactivation of KSHV is associated with major changes in the nuclear organization and redistribution of nucleolar proteins

The lytic cycle of KSHV involves the formation of nuclear replication compartments along with changes in nuclear structures [59]. To characterize these changes, we employed the human renal cell carcinoma SLK cell line and its derivative, iSLK, as model systems [60]. iSLK cells are widely used to study KSHV, as they can stably maintain latent infection and harbor a doxycycline (Dox)-inducible cassette of the KSHV gene product RTA/ORF50, which enables an efficient switch of the lytic cycle when treated with n-Butyrate (n-But). iSLK cells were infected with the recombinant bacterial artificial (BAC) KSHV clone, BAC16, which constitutively expresses GFP from the elongation factor 1-α-promoter [61], or with BAC16-mCherry-ORF45 clone which also expresses the immediate-early lytic gene product ORF45 fused to monomeric Cherry fluorescent protein (mCherry), and thereby allows tracking of cells undergoing lytic replication [62].

To observe the nucleolar organization during the productive cycle of KSHV, we treated BAC16-mCherry-ORF45-infected iSLK cells with Dox and n-But to induce lytic reactivation. Controls included uninfected SLK cells that were similarly treated with Dox and n-But, and uninduced iSLK-infected cells. Of note, as in other experimental cell models of KSHV, the induction of lytic reactivation in infected iSLK cells is asynchronous, and therefore when examining cells post induction we observed different stages of the lytic cycle. These stages can be distinguished by the intensity of the expression and the distribution of lytic viral proteins such as mCherry-ORF45. In addition, late stages of the lytic cycle are characterized by the formation of a large central nuclear zone devoid of chromatin, along with condensed chromatin at the nuclear periphery [9]. Cells were processed 48-hr following lytic induction, by which time a significant proportion of the cells undergo lytic reactivation and DNA replication is well under way; yet, uninduced cells and cells at early stages of the lytic cycle are also evident at this time point. Cells were stained with antibodies to UBF (Fig. 1A), Fibrillarin (Fig. 1B) and Nucleophosmin 1 (NPM1) (Fig. 1C) which are commonly used to mark the FC, DFC and GC nucleolar substructures, respectively. As shown, lytic reactivation, detected through mCherry-ORF45 expression, initially in the nucleus and later in the cytoplasm and close to the plasma membrane, was associated with an extensive destruction of the nuclear architecture and condensation of the nuclear chromatin to the periphery. This was accompanied by changes in the localization pattern of the three tested nucleolar proteins. Of note, since the lytic cycle is asynchronous, images of different stages, characterized by different intensities and cellular distribution of mCherry-ORF45 and by different organization patterns of the chromatin, are presented. The rRNA transcription factor UBF, which is normally associated with rDNA at the rRNA promoters and plays a role in the recruitment of RNA pol I, demonstrated scattered dot-like structures in the nucleoplasm upon lytic induction. The nucleolar protein Fibrillarin, which is the core protein of C/D small nucleolar RNAs (snoRNAs) that functions as 2’-O-methytransferase and is involved both in methylation and processing of rRNA, demonstrated clear changes in its distribution, and was localized in nucleoplasm-dispersed dots along with 1-3 large peripheral foci surrounded by dense chromatin. NPM1, a multifunctional nucleolar protein that normally resides in the GC sub-structure, exhibited two major patterns upon induction of the lytic cycle, characterized either by scattered nucleoplasmic dots along with nucleolar and diffuse nuclear staining, or marginal nuclear staining that overlapped with the chromatin.

**Figure 1.**
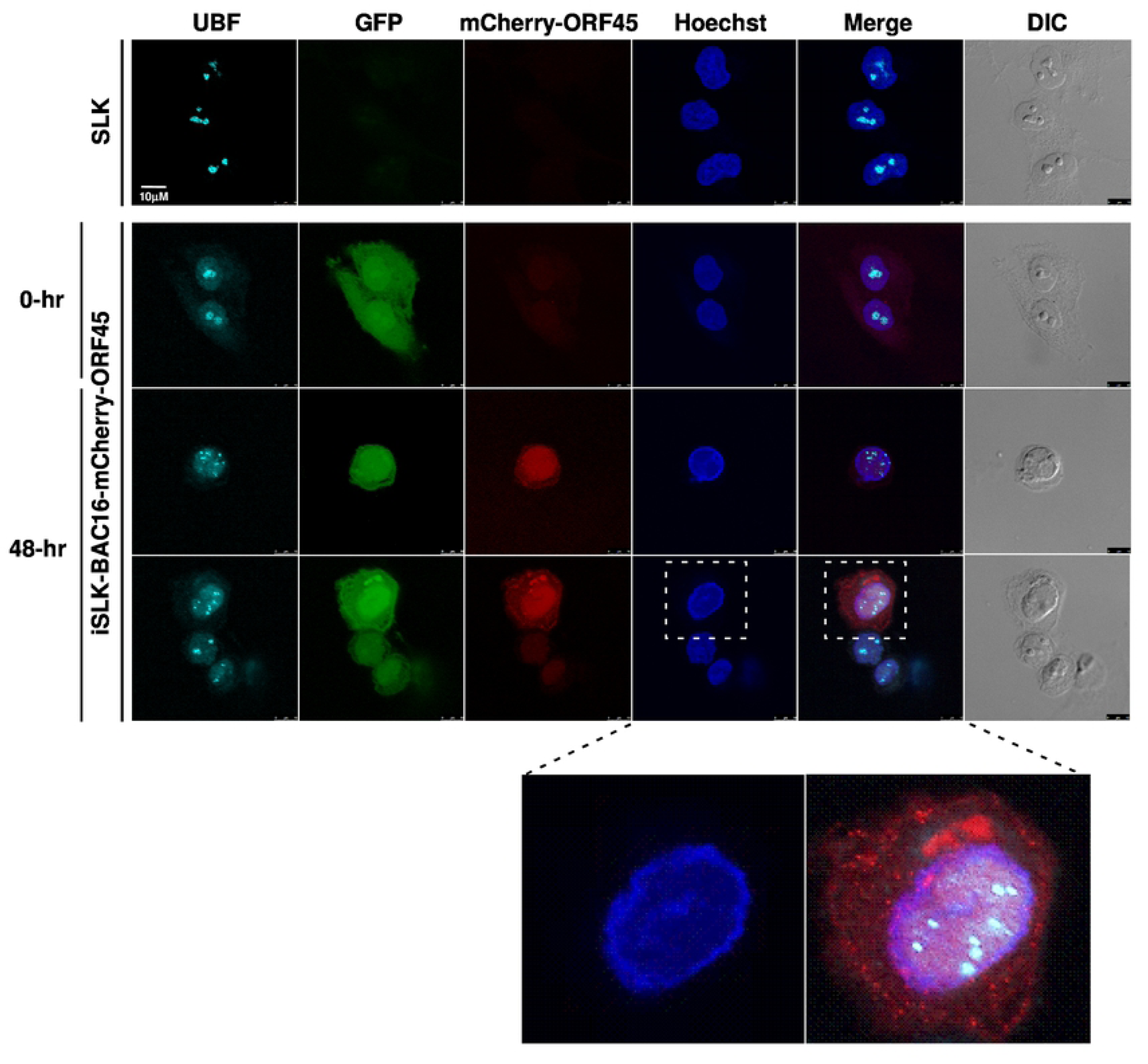
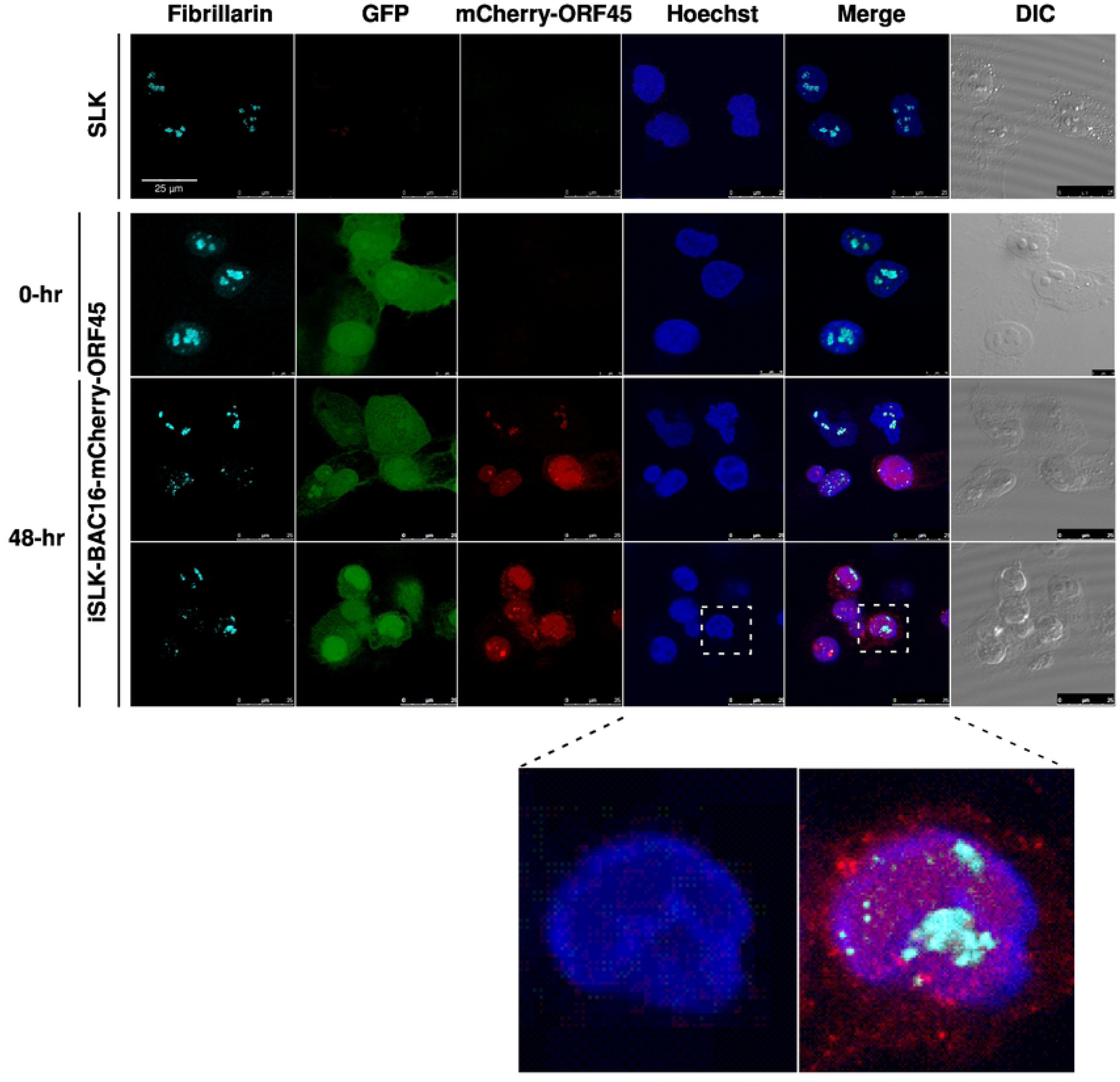
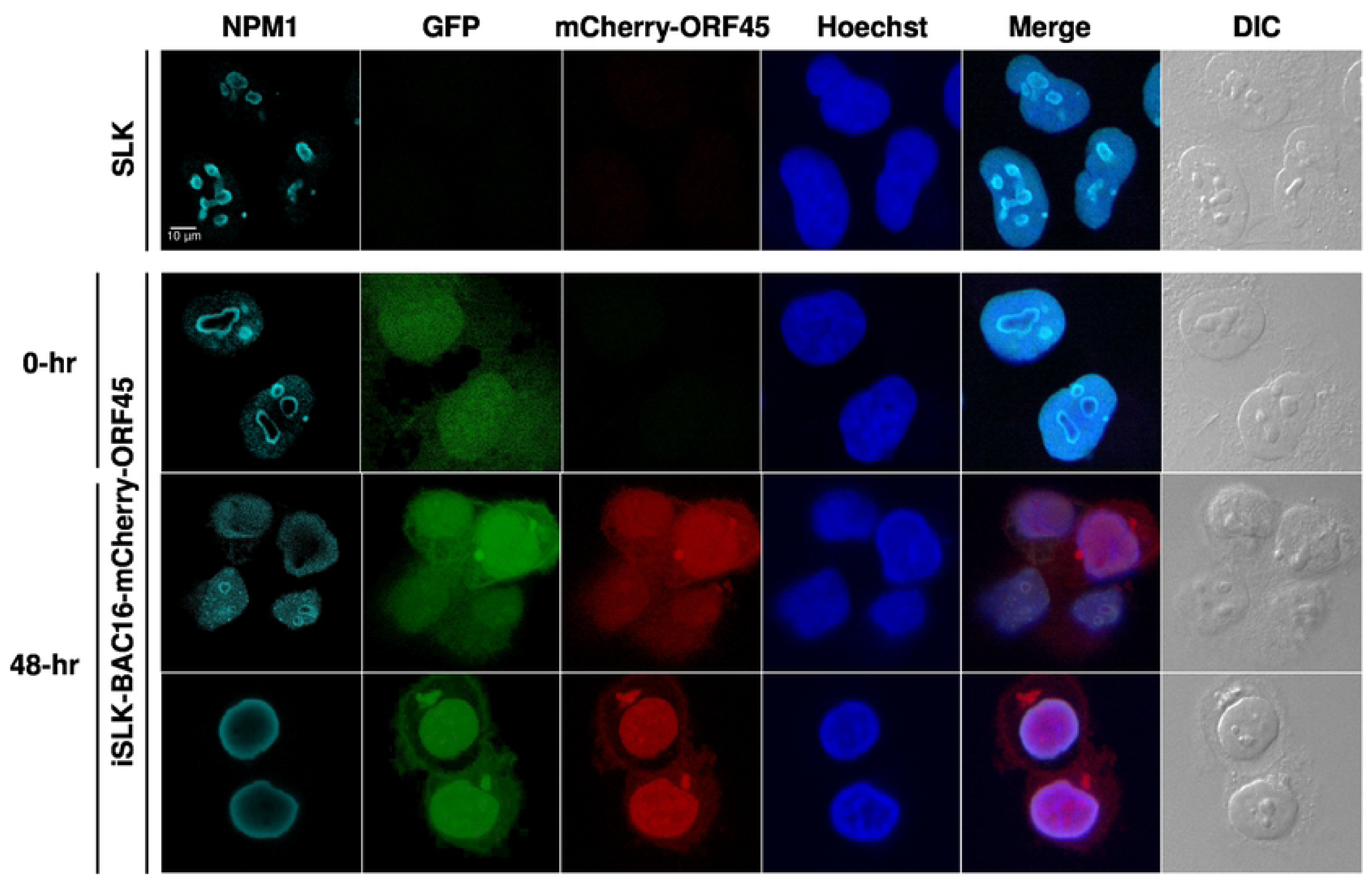
Redistribution of the nucleolar proteins UBF, Fibrillarin and Nucleophosmin 1 (NPM1) upon lytic reactivation of KSHV. BAC16-mCherry-ORF45-infected iSLK cells were treated for 48-hr with 1 µg/ml Dox and 1 mM n-Butyrate to induce viral lytic reactivation. Similarly treated uninfected SLK cells and uninduced infected iSLK cells were used as controls. Cells were stained with antibodies to UBF (A), Fibrillarin (B) or NPM1 (C). Cy5-conjugated mouse secondary antibodies were used for detection. The corresponding staining of nuclear DNA by Hoechst and DIC is also shown. Infected cells expressed GFP, while mCherry-ORF45 expression in iSLK-BAC16-mCherry-ORF45-infected cells marked cells undergoing lytic induction. Cells undergoing lytic reactivation were also identified based on the appearance of reduced chromatin nuclear staining.

### The nucleolar proteins UBF and Fibrillarin do not colocalize with promyelocytic leukemia (PML) nuclear bodies

PML bodies are sub-nuclear structures involved in a wide range of cellular functions, including transcription control, protein modifications, DNA damage response, apoptosis, senescence and antiviral responses [63]. Given the known functions of the PML bodies during viral infections and in view of their well-known dot-like nuclear distribution, we examined potential colocalization of UBF and Fibrillarin with PML bodies during lytic reactivation. As shown in Fig. 2, in line with previous reports [64, 65] a decreased number of PML bodies was evident in cells undergoing lytic replication identified by redistribution of the nucleolar proteins UBF and Fibrillarin. Yet, neither UBF nor Fibrillarin colocalized with foci of PML nuclear bodies during the latent or the lytic cycle of KSHV infection.

**Figure 2.**
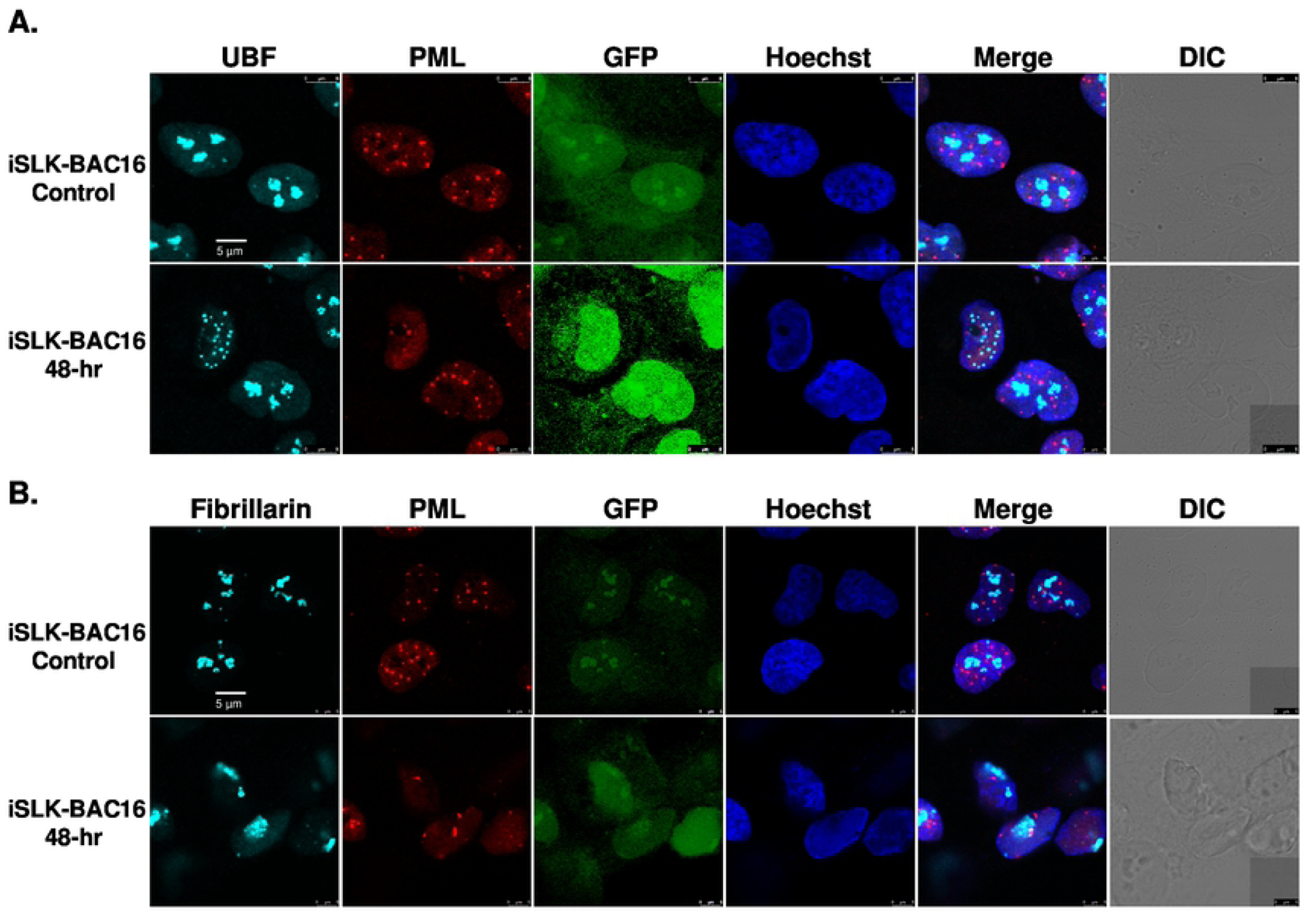
UBF and Fibrillarin do not colocalize with PML bodies. BAC16-infected iSLK cells were treated for 48-hr with 1 µg/ml Dox and 1 mM n-Butyrate to induce lytic reactivation. Uninduced BAC16-infected iSLK cells were used as a control. Cells were stained with anti-PML followed by Rhodamine-conjugated secondary antibody. Subsequently, the cells were incubated with antibodies to UBF (A) or Fibrillarin (B), and then with anti-Rabbit Cy5 secondary antibody. Chromatin was detected by Hoechst staining. Infected cells expressed GFP under the EF1-alpha constitutive promoter.

### The nucleolar protein UBF is partially colocalized with the KSHV latency-associated nuclear antigen 1 (LANA-1)

The latency-associated nuclear antigen 1 (LANA-1) protein, encoded by KSHV, is an essential multifunctional viral protein that controls latency and lytic reactivation, as well as several cellular checkpoints [66, 67]. Given the importance of LANA-1 in KSHV infection and its known nuclear distribution in a speckled pattern, which is reminiscent of the localization of UBF during the lytic cycle, we examined its potential colocalization with UBF. As shown in Fig. 3, UBF was localized in the nucleoli of control latently KSHV-infected iSLK cells, while LANA-1 protein was dispersed in the nucleus in a large number of small foci. At early phases of the lytic cycle (second row), recognized by the partial redistribution of UBF from the nucleoli, a fraction of UBF dots clustered together with LANA-1, though the dots did not colocalize. Partial colocalization of UBF and LANA-1 was evident at later stages of the lytic cycle (bottom row), as evident by complete dispersion of UBF into small dots and the appearance of a chromatin-depleted nucleus.

**Figure 3.**
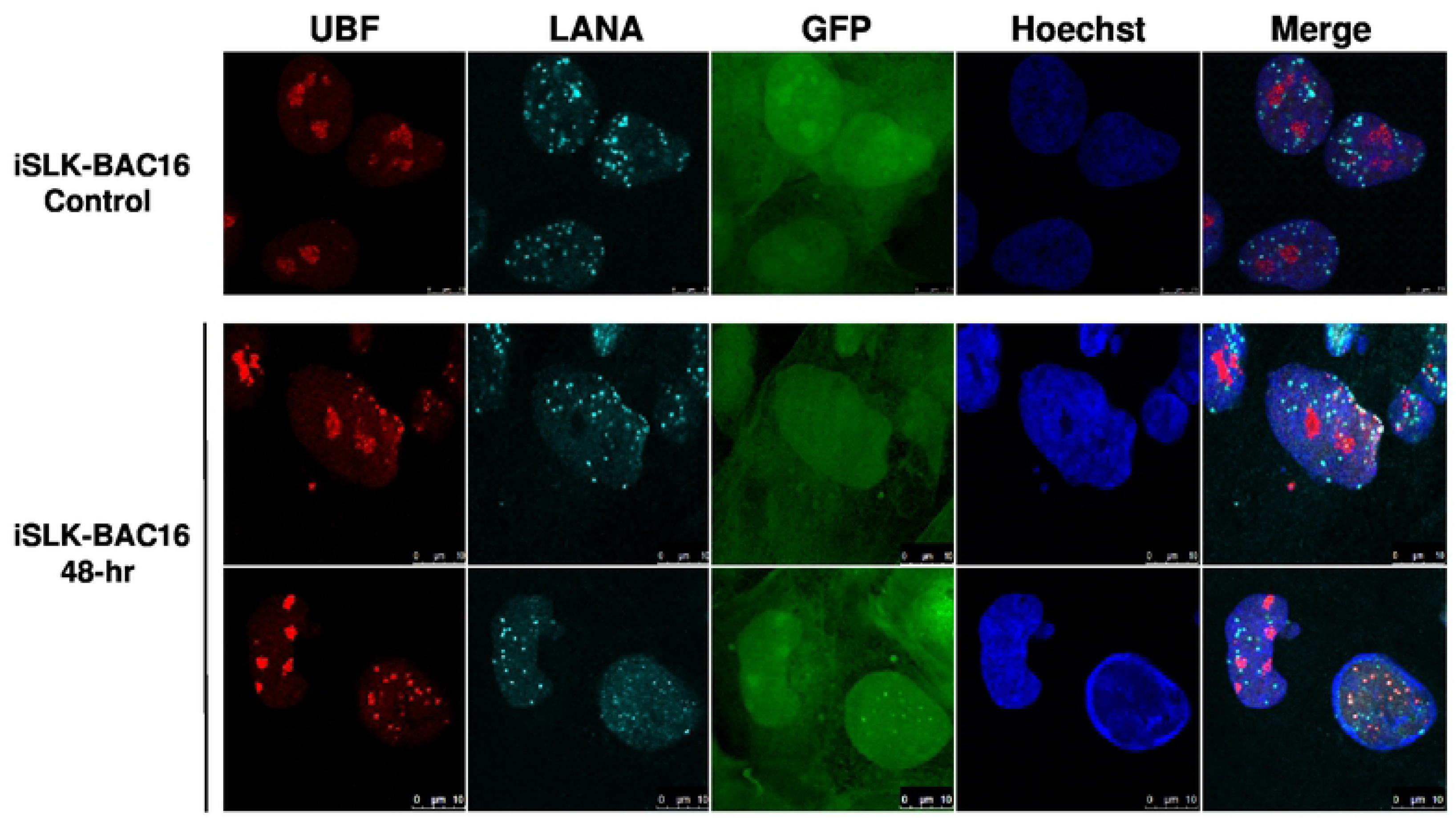
Cellular distribution of UBF and LANA-1 proteins during latency and lytic reactivation of KSHV. BAC16-infected iSLK cells were treated for 48-hr with 1µg/ml Dox and 1 mM n-Butyrate to induce lytic reactivation. Uninduced BAC16-infected iSLK cells were used as a control. Cells were stained with anti-LANA-1 followed by Cy5-conjugated anti-rat secondary antibody. Subsequently, the cells were incubated with antibodies to UBF and then with anti-rabbit Cy5-conjugated secondary antibody. Chromatin was detected by Hoechst staining, and viral infection was confirmed by GFP expression.

### Redistribution of UBF and Fibrillarin does not depend on viral DNA replication or late viral gene expression

Redistribution of Nucleolin and NPM1 during HSV-1 infection was previously shown to depend on the expression of the late viral protein UL24 [42-44, 68]. Accordingly, we examined whether treatment with the viral DNA polymerase inhibitor phosphonoacetic acid (PAA), which inhibits virus replication and expression of late viral genes, would alter the redistribution of UBF and Fibrillarin during lytic reactivation of KSHV. As shown in Fig. 4, treatment with PAA did not affect the relocalization of either UBF or Fibrillarin. This implies that the redistribution of these proteins during lytic reactivation does not depend on viral DNA replication or on the formation of viral replication compartments, nor on late viral genes that are not actively expressed under PAA treatment.

**Figure 4.**
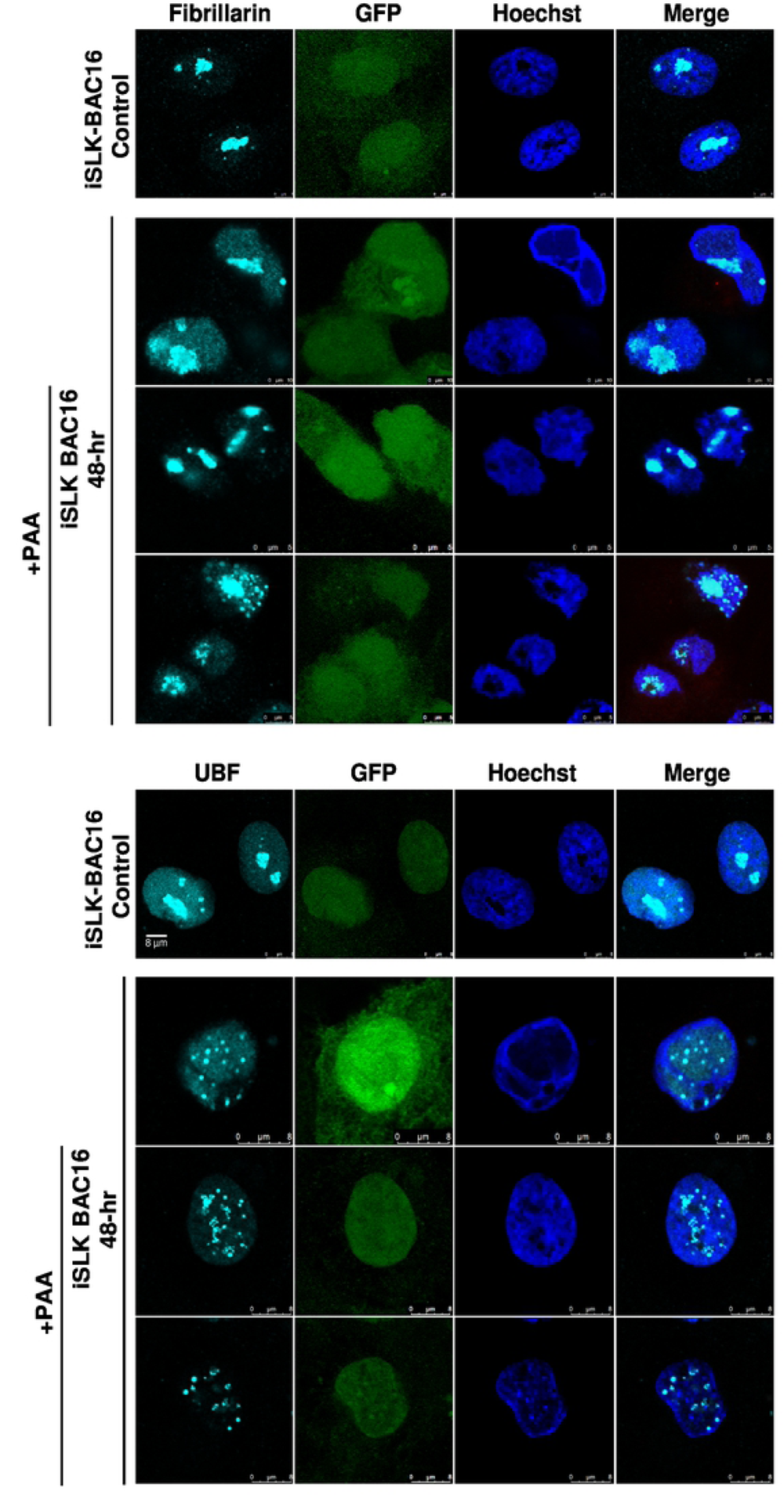
Redistribution of UBF and Fibrillarin does not depend on viral DNA replication. BAC16-infected iSLK cells were treated for 48-hr with 1µg/ml Dox and 1 mM n-Butyrate to induce lytic reactivation. Treatment with PAA was carried 2-hr prior to lytic induction and continued during lytic reactivation. BAC16-infected iSLK cells that were either left untreated or induced to undergo lytic reactivation with no PAA treatment were used as control. Cells were stained with anti-UBF (A), or anti-Fibrillarin (B) and then incubated with anti-mouse Cy5-conjugated secondary antibody. Chromatin was detected by Hoechst staining. Infected cells expressed GFP under the EF1-alpha constitutive promoter.

### UBF is partially and transiently colocalized with replication compartments, whereas Fibrillarin does not colocalize with replication compartments during lytic induction

Next, we examined the cellular localization of UBF and Fibrillarin along with ORF59 viral protein which functions as a processivity factor during viral DNA replication, and hence is localized in the viral replication compartments during the lytic cycle [69-71]. As shown in Fig. 5, and Movies S1 and S2, UBF began to disperse early during lytic reactivation, when expression of ORF59 was undetectable. Later, during lytic reactivation, UBF was clearly observed within the replication compartments, and a small number of UBF foci overlapped with ORF59. At late stages of reactivation, when ORF59 was visible at the nuclear margins, no colocalization was observed between the two. This finding suggests that UBF transiently colocalizes with replication compartments at early phases of the lytic cycle. In contrast, Fibrillarin did not share common localization with ORF59. Rather, at early phases of the lytic cycle, when Fibrillarin began to redistribute, ORF59 occupied most of the nuclear zone, whereas at later times, when Fibrillarin appeared in large structures at the nuclear margins, ORF59 formed a ring-like structure at the periphery of the cell nucleus. Finally, Nucleolin, which was previously reported to participate in the replication of HSV-1 and hCMV [40, 72, 73], also redistributed to the periphery of the cell nucleus while partially remaining in large structures, colocalized with ORF59 at the nuclear periphery.

**Figure 5.**
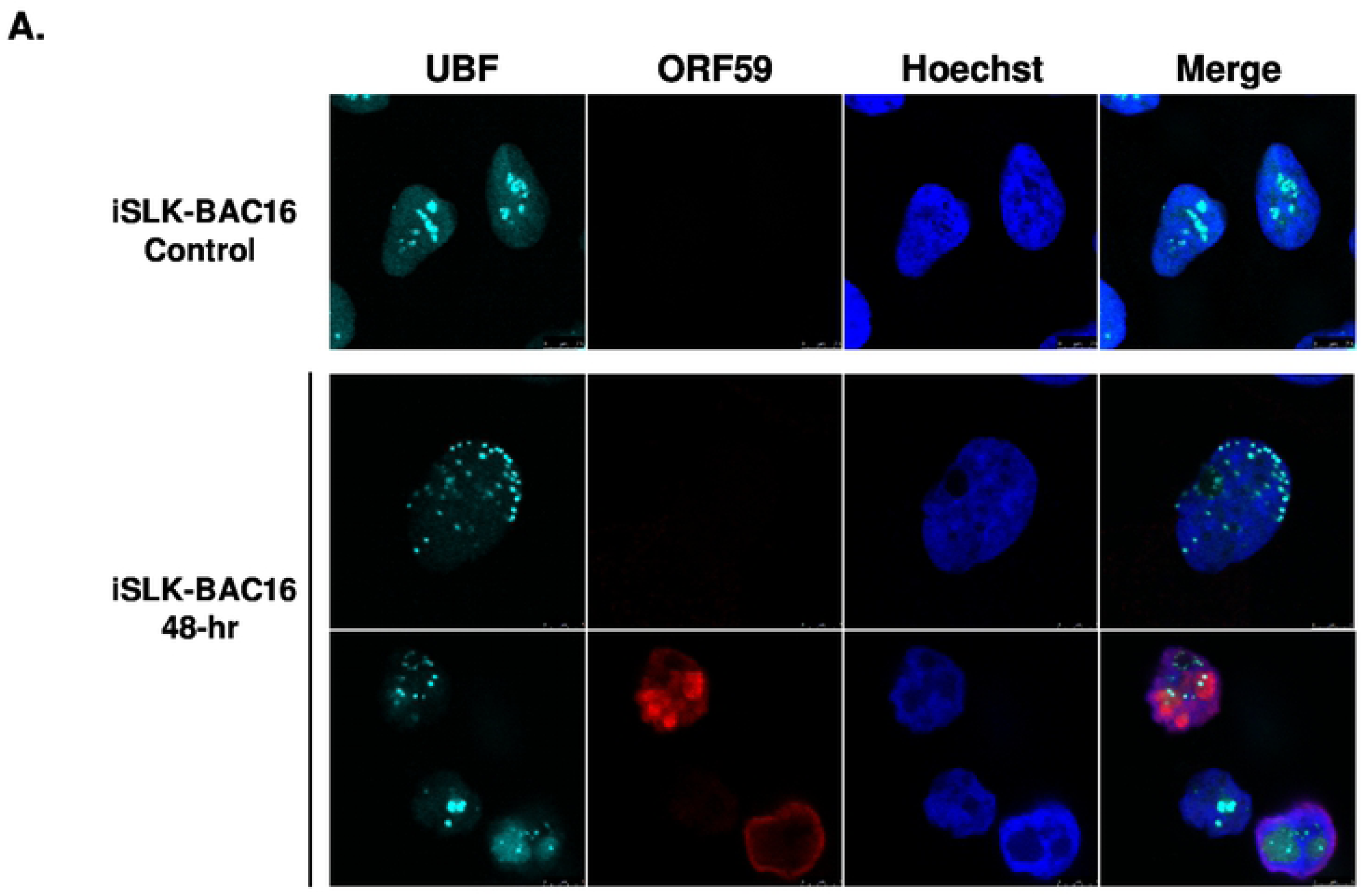
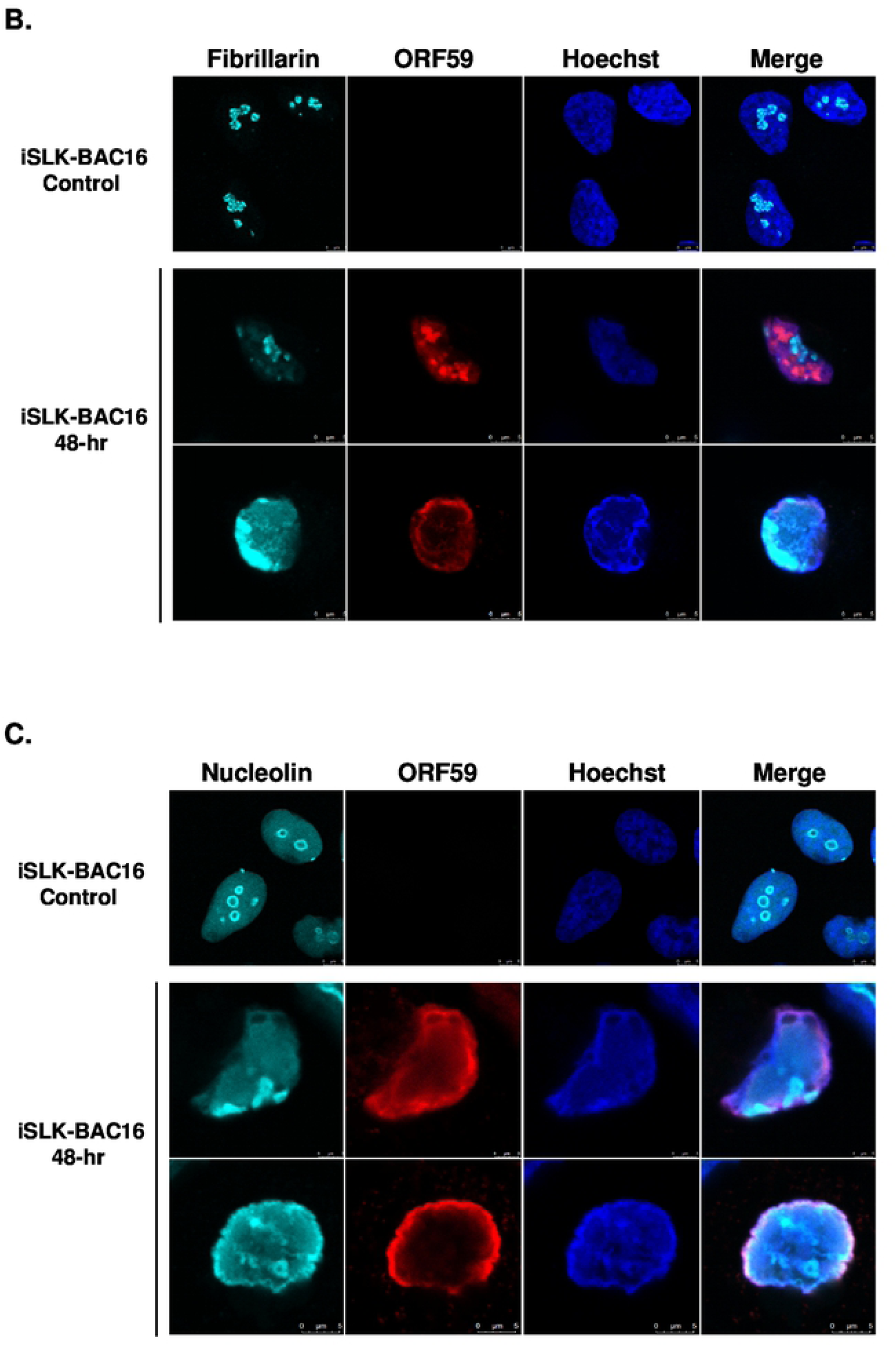
UBF partially and transiently colocalizes with viral replication compartments whereas Fibrillarin does not colocalize with replication compartments during lytic reactivation. BAC16-infected iSLK cells were treated for 48-hr with 1µg/ml Dox and 1 mM n-Butyrate to induce lytic reactivation. Untreated infected iSLK cells were used as control. Cells were stained with anti-ORF59 and secondary Rhodamine-conjugated antibody. Cells were subsequently incubated with anti-UBF (A), anti-Fibrillarin (B), or anti-Nucleolin (C) and then incubated with anti-Rabbit Cy5-conjugated secondary antibody. Chromatin was detected by Hoechst staining.

### Redistribution pattern of UBF during lytic reactivation overlaps with the nucleolar proteins RPA194 and PICT-1 but not with Fibrillarin

As UBF and Fibrillarin appear to exhibit different redistribution patterns and kinetics after lytic reactivation, we next examined their cellular localization at the same cells and with additional nucleolar proteins as well. As shown in Fig. 6A and Movie S3, UBF and Fibrillarin demonstrated different dispersion patterns with respect to each other and their dispersion kinetics were also different. At early activation stages, when the chromatin distribution was still normal, UBF already began to disperse, though most of it was still found in the nucleoli. At this stage, Fibrillarin did not appear to be scattered at all, and remained within large nuclear foci that likely represented the nucleoli (Fig. 6A, second row). At more advanced lytic reactivation stages, there was little overlap between UBF, which appeared to be scattered throughout the nucleoplasm, and Fibrillarin which was found mainly in the nuclear periphery. In contrast, UBF and the large RNA polymerase I subunit RPA194 shared similar redistribution kinetics and pattern after lytic reactivation in all cells examined (Fig. 6B). Similarly, the nucleolar exosome recruiting and assembly factor, PICT-1 [74, 75] also shared redistribution kinetics and pattern with UBF (Fig. 6C). Finally, partial colocalization was evident between UBF, which formed small dots, and Nucleolin, which redistributed to the nucleoplasm, probably within replication compartments (Movie S4). These findings suggest that the dispersion of nucleolar proteins during lytic reactivation is regulated. Proteins that are likely to share common roles remain co-localized, whereas other proteins disperse to different locations and hence may have distinct functions. Furthermore, the common cellular distribution of the rRNA transcription factor UBF, the RNA polymerase I subunit RPA194, and PICT-1 suggests that rRNA transcription in these cells is maintained.

**Figure 6.**
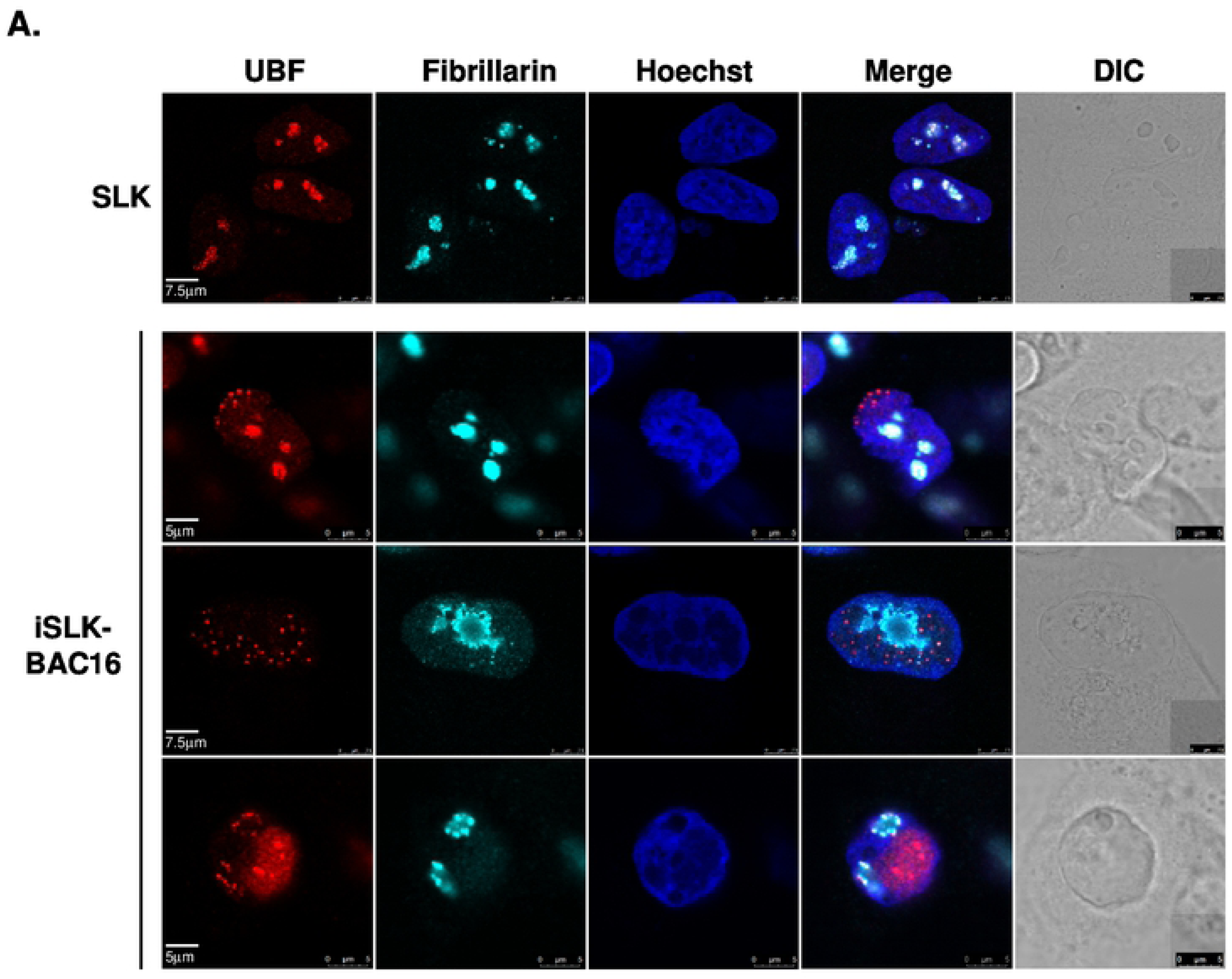
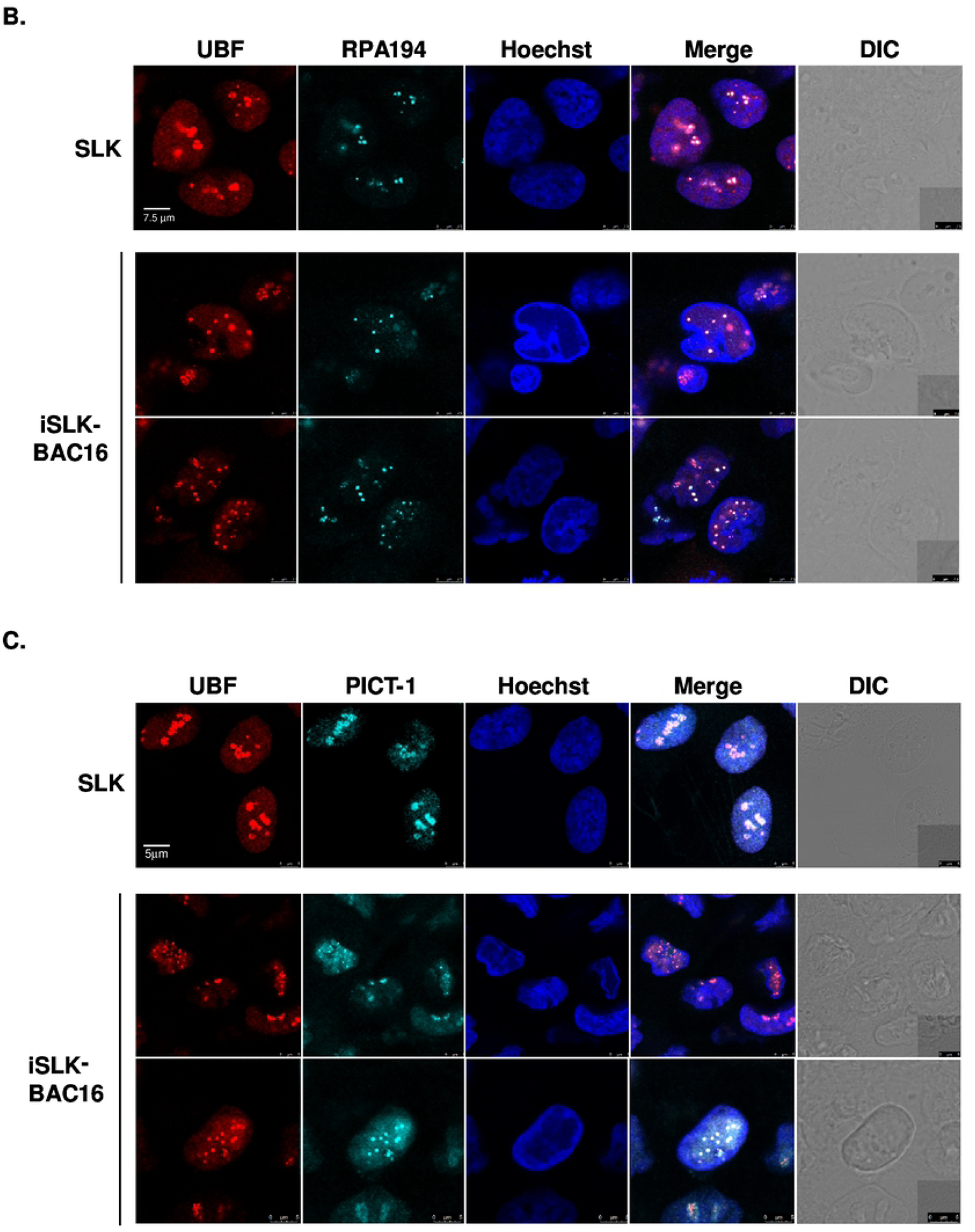
UBF shares common redistribution kinetics and pattern with RPA195 and PICT-1 but not with Fibrillarin during lytic reactivation. BAC16-infected iSLK cells were treated for 48-hr with 1µg/ml Dox and 1 mM n-Butyrate to induce lytic reactivation. Uninfected SLK cells were similarly treated and used as control. Cells were stained with mouse anti-UBF and Rhodamine-conjugated secondary antibodies. Cells were then incubated with anti-Fibrillarin, which was detected by anti-rabbit Cy5-conjugated secondary antibody (A). Cells were stained with rabbit anti-UBF and Cy3-conjugated secondary antibodies, and then with anti-RPA194 (B) or anti-PICT-1 (C), which were detected by anti-mouse Cy5-conjugated secondary antibody. Chromatin was detected by Hoechst staining.

### rRNA transcription and processing persist during lytic reactivation of KSHV

In mammalian cells, the initial transcription product of rDNA is known as the 47S pre-rRNA. Processing of this transcript involves a series of cleavages and chemical modifications and results in the production of intermediate rRNAs, including 45S, 41S and 32S rRNA, and the three mature rRNAs: the 28S, 18S and 5.8S species. Given the changes in the nuclear organization and in the distribution of key nucleolar proteins, we aimed to characterize changes in the synthesis, localization and processing of the rRNA. First, we quantified pre-rRNA levels using RT-qPCR with primers that target the 5’-external transcribed spacer (5’-ETS) which is included in the 47S pre-rRNA transcript but not in the mature rRNA. As shown in Fig. 7, no significant changes in the levels of pre-rRNA were detected upon lytic induction, suggesting that transcription of pre-rRNA continues during productive KSHV infection.

**Figure 7.**
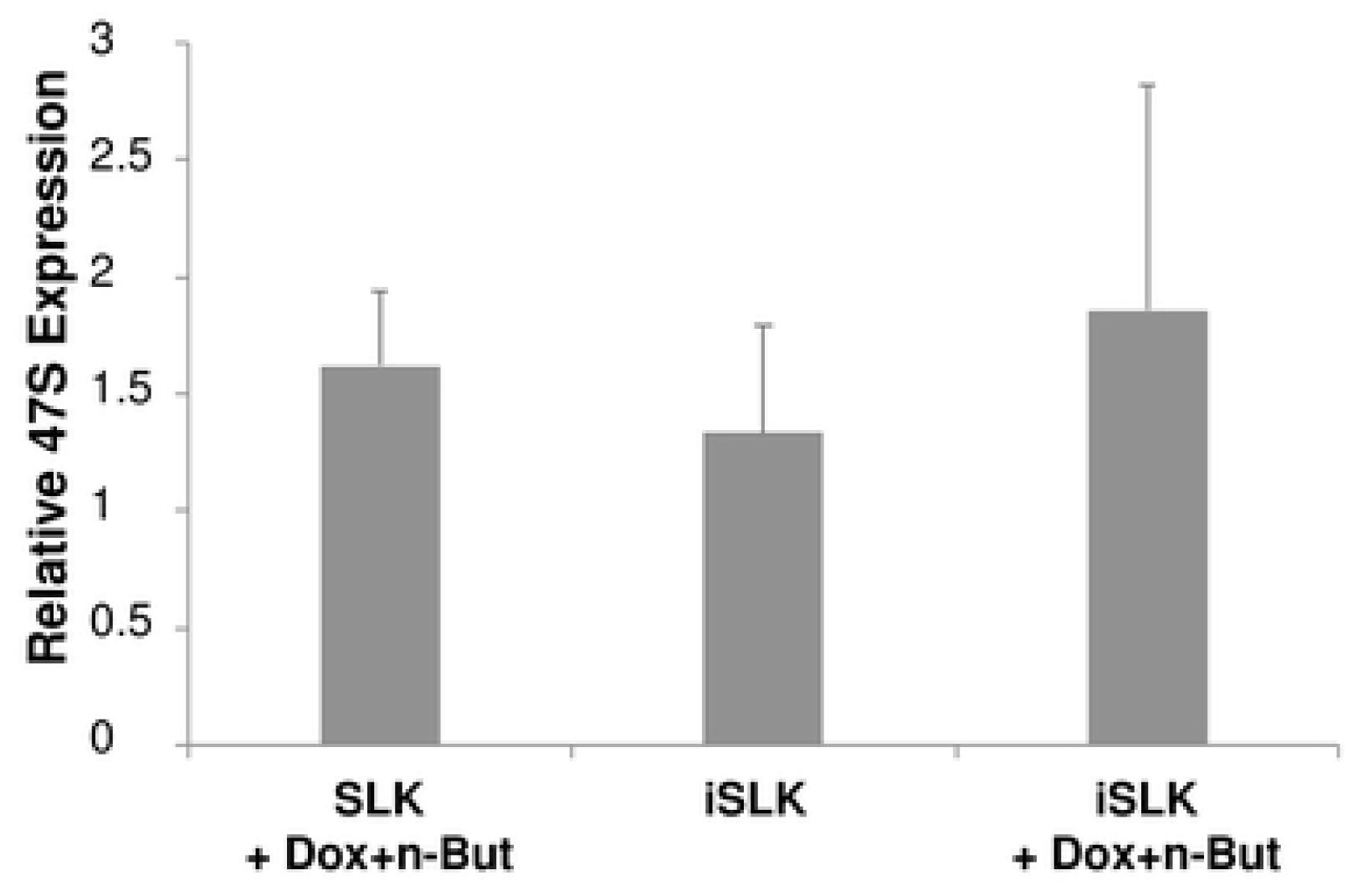
Lytic reactivation does not alter expression levels of the 47S pre-rRNA. Uninfected SLK and BAC16-mCherry-ORF45-infected iSLK cells were treated for 48-hr with 1µg/ml Dox and 1 mM n-Butyrate to induce lytic reactivation. Untreated cells were used as controls. Lytic reactivation was confirmed by expression of mCherry-ORF45 in ∼80% of cells. 47S pre-rRNA levels were determined using RT-qPCR with primers that target the 5’ external transcribed spacer (ETS) of pre-rRNA, and the results obtained were normalized to 18S rRNA. Data are presented relative to untreated SLK cells, and are from averages of five biological repeats, each including three technical replicates. No statistically significant difference between SLK and iSLK cells prior or after lytic reactivation were obtained (one-way ANOVA and One-sample T-test).

As in most other cell culture models for KSHV infection, the lytic reactivation in iSLK cells is not synchronized. Hence, it was important to characterize the rRNA transcription kinetics with additional assays focusing on individual cells. To this end, we performed a fluorescent in-situ hybridization (FISH) assay in which we labeled newly synthesized rRNA transcripts with the UTP analog BrU within a limited time frame of 30 min. Lytic reactivation was identified through the redistribution of UBF. As shown in Fig. 8, a large fraction of the BrU signal was not nuclear, probably due to limited penetration of the analog into the nucleus or as a result of export of newly synthesized ribosomes. Yet, clear large foci of BrU staining, which overlapped with UBF staining were evident in control SLK cells. A large number of small BrU foci, representing newly synthesized rRNA transcripts, were detected within the nucleus during lytic reactivation when UBF formed small nuclear dots. Some rRNA foci were noted adjacent to UBF. The incorporation of BrU clearly indicates that rRNA synthesis persists during lytic reactivation. In addition, no accumulation of selected pre-rRNA intermediates was evident by Northern blot analysis, suggesting that proper processing of pre-rRNA continued (Fig. 9). Of note, unique rRNA processing products were previously identified in HSV-1 infected cells during lytic infection [47]. Yet, no unique products were identified here, and no significant differences in the relative expression levels of the different processing products were detected. As the nucleolar protein Fibrillarin is known to participate in the processing of pre-rRNA, we examined the cellular localization of this protein along with the relatively long-lived internal transcribed spacer 1 (ITS-1) pre-rRNA sequence by using FISH combined with immunofluorescence. As shown in Fig. 10, in SLK control cells, the ITS-1 fluorescent signal accumulated mainly in nucleoli and colocalized with Fibrillarin. However, during lytic reactivation, identified through the marginalization of the nucleolar structure and by low-density chromatin, the ITS-1 was detected as dots, and mostly overlapped with Fibrillarin. This finding further indicates that the processing of rRNA continues during lytic reactivation.

**Figure 8.**
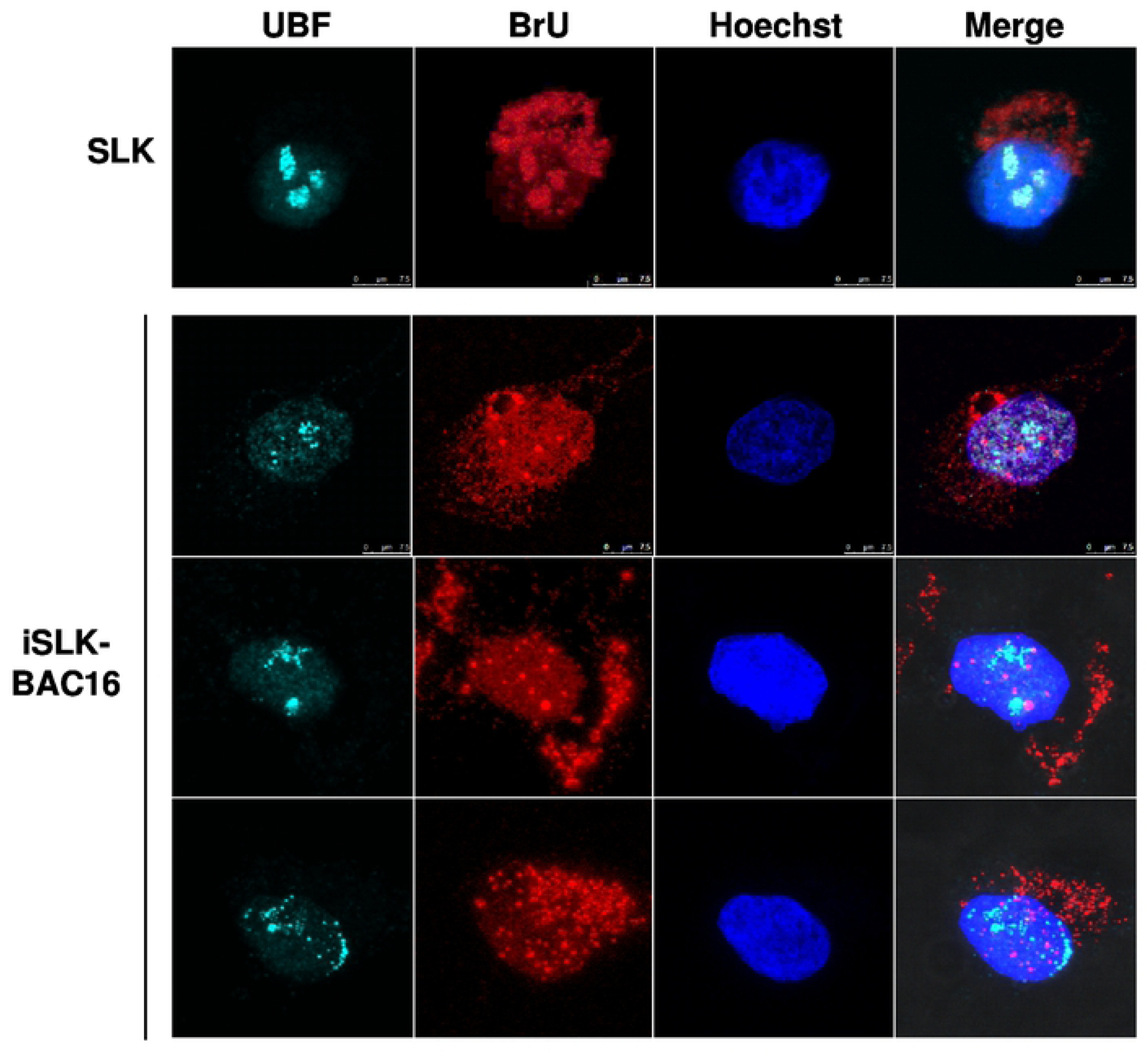
FISH analysis of newly synthesized rRNA transcripts in cells during viral lytic reactivation. BAC16-infected iSLK cells were treated with 1µg/ml Dox and 1 mM n-Butyrate to induce lytic reactivation. Uninfected SLK cells were used as control. After 48-hr, cells were permeabilized with lysolecithin and incubated for 30 min with labeling buffer containing BrU and alpha-amanitin to block RNA polymerase II transcription. Cells were then fixed and stained with BrU antibody and secondary anti-mouse Rhodamine-conjugated antibody. Cells were then stained with rabbit anti-UBF and Cy5-conjugated anti-rabbit secondary antibody. Hoechst staining was used to detect chromatin.

**Figure 9.**
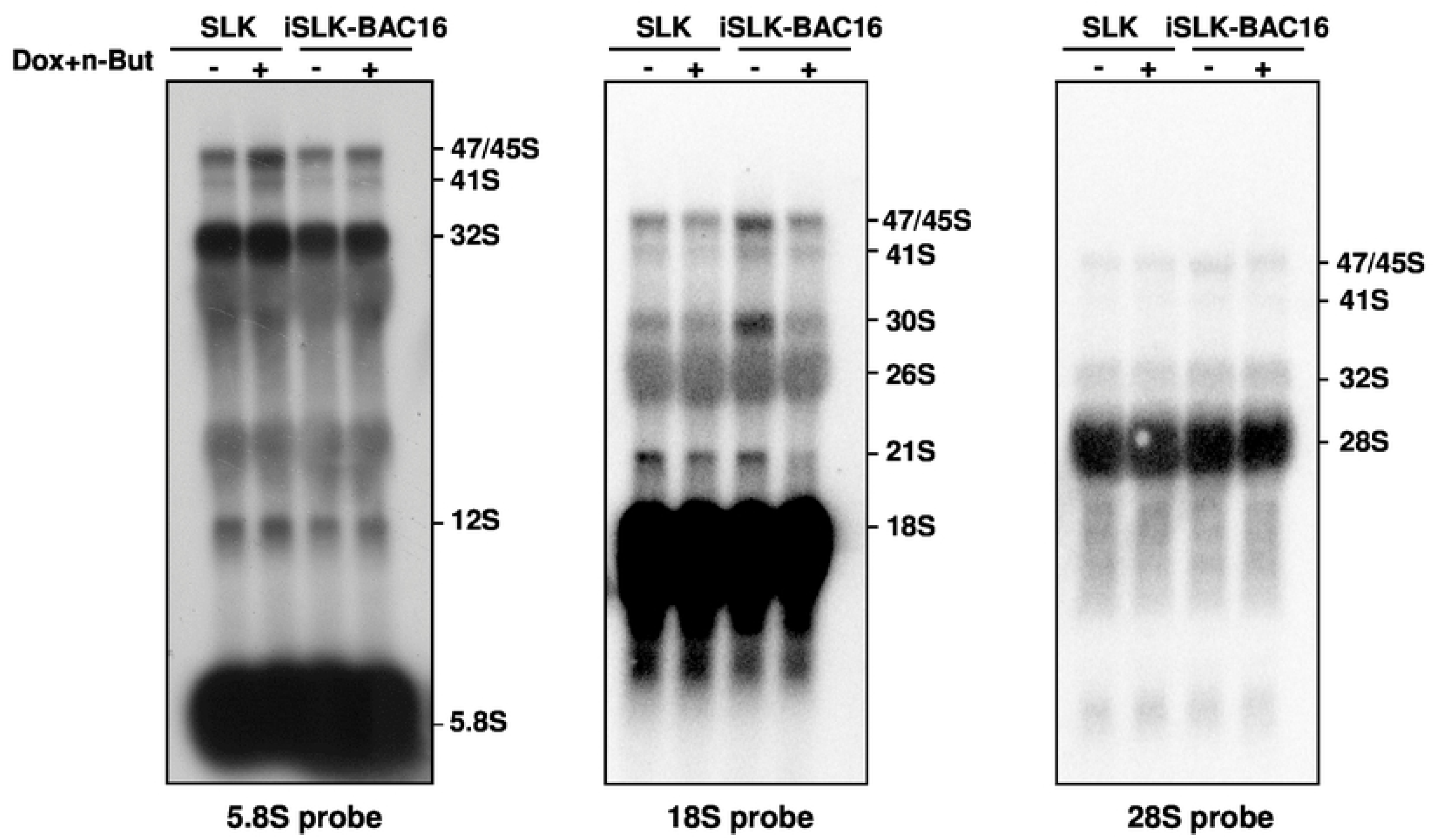
Northern blot analysis of rRNA during viral lytic reactivation. Total RNA was extracted from uninfected SLK and BAC16-mCherry-ORF45-infected iSLK cells that were either left untreated or treated with Dox and n-Butyrate for 48-hr. Lytic reactivation was confirmed by expression of mCherry-ORF45 in ∼80% of cells. 20 µg RNA from each sample was resolved on denaturing 1.2% agarose gel, transferred to nylon membranes, and hybridized with probes representing 5.8S, 18S and 28S rRNA. No accumulation of pre-rRNA processing products was evident upon lytic reactivation.

**Figure 10.**
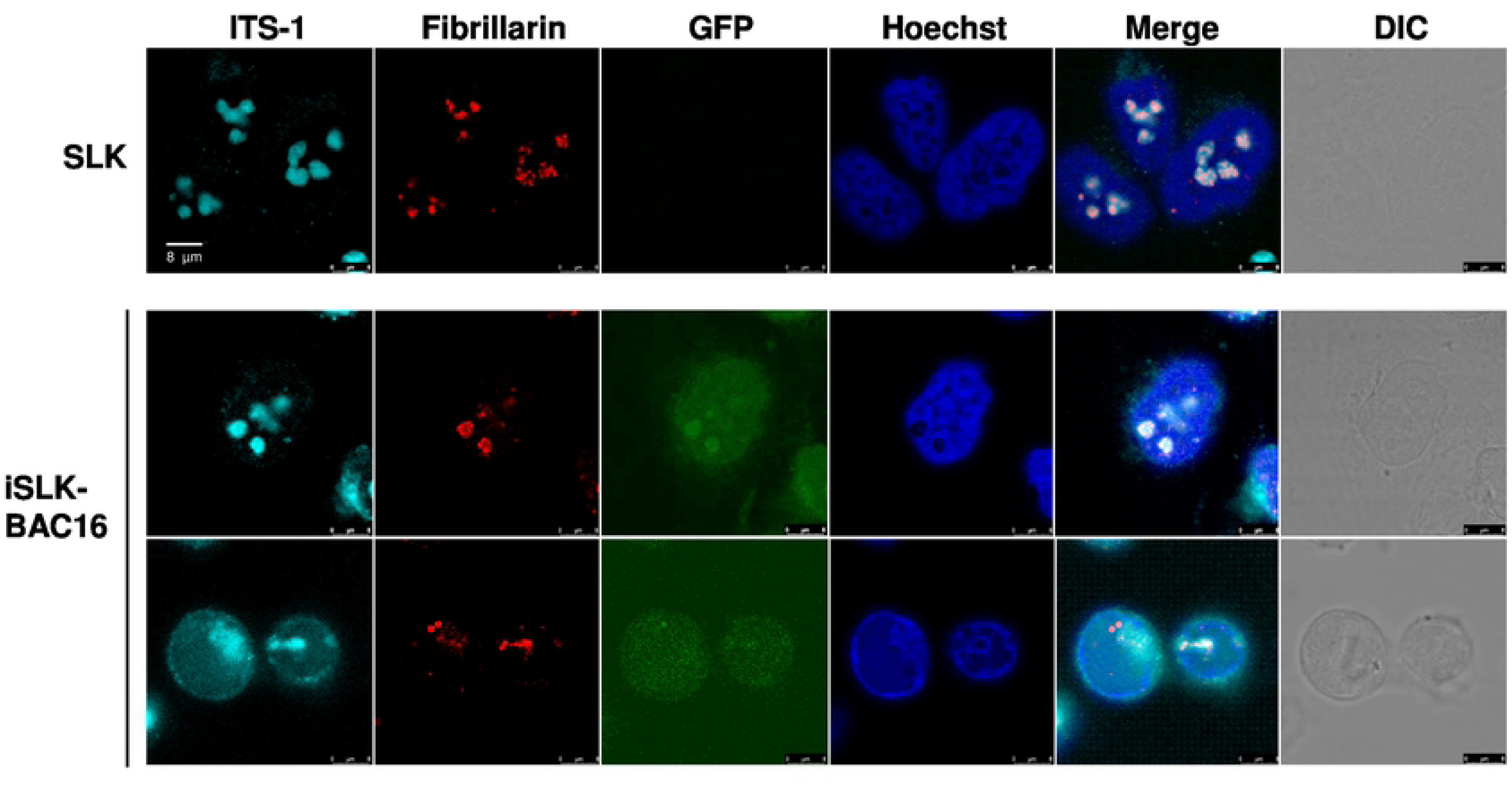
RNA *in situ* hybridization analysis to determine the location of the pre-rRNA transcripts and Fibrillarin in cells during viral lytic reactivation. SLK and BAC16-infected iSLK cells were treated with 1µg/ml Dox and 1 mM n-Butyrate. Cells were incubated with ITS-1 probe and stained with anti-Fibrillarin and Cy3-conjugated secondary antibodies. Hoechst staining was used to detect chromatin.

Taken together, our results suggest that rRNA transcription and processing proceed properly during lytic reactivation of KSHV, despite the different distribution patterns of UBF and Fibrillarin, which are known to participate in these processes. However, while under normal conditions these processes are coupled, under lytic reactivation, the processes can be uncoupled, yet remain functional.

### Changes in pseudouridine levels at specific rRNA sites

Recently, it was postulated that rRNA modifications may provide an important source of ribosome heterogeneity, and tune ribosome function in response to various signals [76-81]. These modifications, which may be fractional at certain positions, stabilize the secondary and tertiary structure of the rRNA scaffold, thereby modulating the efficiency and accuracy of protein translation. One of the most abundant rRNA modifications in eukaryotes involves isomerization of uridine to pseudouridine (Ψ). To identify Ψ sites which change during lytic reactivation, we mapped the Ψ across the rRNA based upon CMC (N-cyclohexyl-N’-β-(4-methylmorpholinium) ethylcarbodiimide p-tosylate) modification followed by alkaline treatment. Under these conditions, the reaction of Ψ sites with CMC results in inefficient reverse transcription during the library preparation process, with the reverse transcription product terminating one nucleotide before the modified base [82]. We prepared RNA-seq libraries from total RNA from BAC16-infected iSLK and control SLK cells that were treated with Dox and n-But to induce lytic reactivation with and without CMC treatment. The ratio of the number of reads supporting reverse transcriptase termination to the number of reads overlapping it (known as the Ψ-ratio) was calculated [83]. The Ψ-fold change (Ψ-fc) is the log2-transformed Ψ-ratio of the treated samples (+CMC) divided by the Ψ-ratio in the non-treated ones (−CMC). Comparing the Ψ-fc across replicates identified changes in Ψ levels at selected positions, illustrated in Fig. 11A and detailed in Supplementary Table T1. The most conspicuous changes observed were the increase in modification (hyper-modifications) at six positions - two located in domain IV (H69; LSU_ Ψ3695 and Ψ3758), one in domain VI (LSU_ Ψ4973), and two in or around the peptidyl transferase center (PTC; LSU_ Ψ4361 and Ψ4457). Hypomodified positions were identified in domain VI (LSU_ Ψ4636 and Ψ4689) and one in the PTC (LSU_ Ψ4521) (Fig. 11B and Supplementary Fig. S1). Selected changes were further verified by primer extension (Fig. 11C). These results are reminiscent of changes observed in cancer [84], whereby increases and decreases in distinct Ψ positions were observed. Note that the changes take place in functionally important domains of the rRNA and therefore may affect translation.

**Figure 11.**
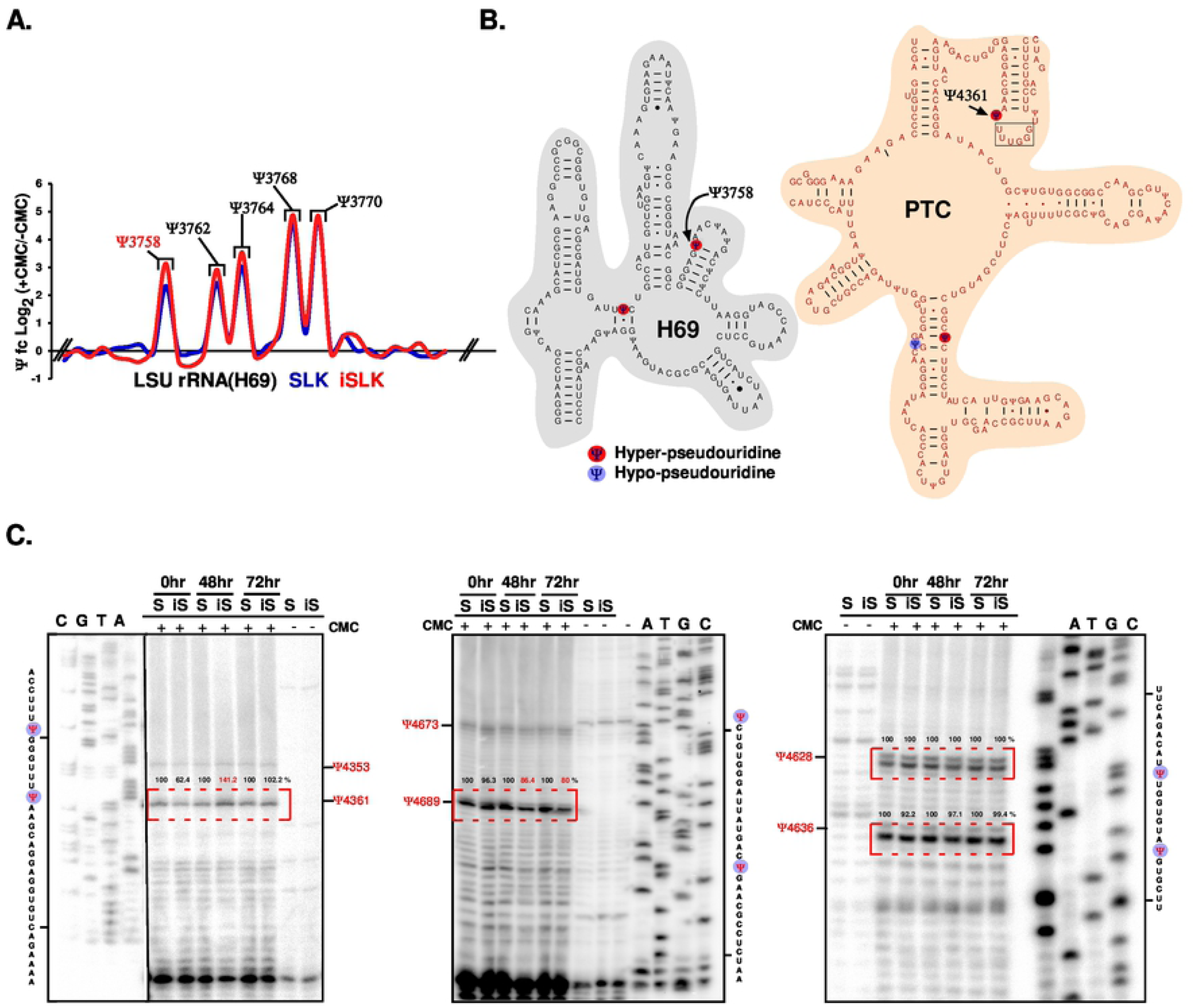
Changes in Ψ levels detected in KSHV-infected iSLK cells undergoing lytic reactivation. Uninfected SLK cells and BAC16-infected iSLK cells were treated with Dox and n-Butyrate for 48-hr. RNA-seq libraries from nuclear RNA were prepared with or without CMC (N-cyclohexyl-N9-b-(4-methylmorpholinium) ethylcarbodiimide p-tosylate). A representative line-graph of the Ψ-fc(log2) of BAC16-infected iSLK and uninfected SLK cells, representing Ψ-ratio of CMC-treated (+CMC) divided by Ψ-ratio of untreated samples (-CMC), is presented for LSU rRNA (H69 domain (A). Scheme representing the position of hyper/hypo-modified Ψ on the secondary structure of rRNA based on Ψ-seq, highlighting functional domains (H69 and PTC) of the rRNA. The structure of human rRNA was derived from (http://www.rna.ccbb.utexas.edu/) (B). Ψ verification by primer extension. Nuclear RNA, treated with CMC (+) or untreated (-), was subjected to primer extension with region-specific primers, and analyzed on 12% denaturing polyacrylamide gel. The results, along with DNA sequencing, are presented for uninfected SLK (S) and BAC16-infected iSLK (iS) cells that were treated with Dox and n-Butyrate for the indicated time points. The positions of Ψ are indicated. As shown, increased pseudouridine level at position U4361, and decreased pseudouridine level at position U4689 was confirmed (C).

## Discussion

The protein content and organization of the nucleolus are dynamic and change under different stress conditions, such as transcriptional inhibition and DNA damage [85]. Major alterations in the morphology and composition of the nucleoli have been previously reported following HSV-1 infection [40, 42, 43]. Nevertheless, persistence of rRNA transcription and ribosome biogenesis, involving modified rRNA maturation, through late stages of HSV-1 infection has been documented [45, 47]. However, almost no description of the nuclear and the nucleolar morphology during KSHV infection is available to date.

By using immunofluorescence microscopy, we documented an extensive and progressive destruction of the nuclear architecture during lytic reactivation of KSHV, with significant condensation of the nuclear chromatin to the periphery of the nucleus. This was associated with extensive redistribution of the nucleolar proteins UBF, Fibrillarin and NPM1 that normally reside at three distinct sub-nucleolar sites. The RNA polymerase I transcription factor UBF, the large RNA polymerase I subunit RPA194, and the vBcl-2 nucleolar partner PICT-1 [57], which also functions in ribosomal biogenesis and in attenuation of the interferon response [75, 86], concomitantly redistributed and colocalized in dot-like structures. The RNA methyltransferase Fibrillarin redistributed later, presenting a few large foci surrounded by dense chromatin, and only partially colocalized with UBF. NPM1 was dispersed in a diffuse nucleoplasmic pattern, and Nucleolin was mostly found in RCs and at the periphery of the cell nucleus. These findings are in line with reported observations during HSV-1 infection [43]. Similarly, Nucleolin was found in the exterior of hCMV replication compartments and was suggested to be important for proper nuclear and subnuclear localization of replication compartment protein [72, 73]. Of note, neither UBF nor Fibrillarin colocalized with PML bodies or with LANA-1 dots, and redistribution of these proteins was not dependent on viral DNA replication and late gene expression.

These experiments suggest that nucleolar proteins undergo nonrandom programmed dispersion that maintains certain protein complexes, but disrupts others during productive infection. Furthermore, it is possible that the same mechanisms and functions engage the localization of UBF, RPA194 and PICT-1, which may participate in related processes during infection. It is not currently known whether these proteins acquire additional functions during lytic reactivation, and whether they continue to function according to their known roles. In addition, it is not known whether the dispersion represents a cellular response to viral infection, or whether it take place due to the formation of replication compartments that generate condensed peripheral chromatin and thus may favor virus replication. Nevertheless, recruitment of particular nucleolar proteins to replication compartments, such as Nucleolin, which was previously reported to participate in the formation of adenovirus and hCMV replication compartments [39, 72, 73], may indicate a role for these proteins in virus replication and/or assembly. Of note, analysis of the proteome associated with adenovirus and HSV-1 genomes illustrated enrichment of nucleolar proteins, including components of RNA polymerase I and the processome small subunit complex, which were suggested to promote expression of viral genes [48]. Whether lytic replication and the changes in the nucleolar architecture observed during the lytic reactivation of KSHV stimulate p53-dependent or independent stress responses, which in turn may affect cell cycle progression and host metabolic pathways, is currently unknown.

Why viral proteins localize to the nucleolus and how nucleolar perturbations affect virus infection have not been precisely defined to date. It is likely that viruses utilize the pluri-functional nature of the nucleolus to enhance virus replication or to avoid cell surveillance mechanisms; yet, it is also possible that nucleolar alterations signify cellular anti-viral responses. It has also been postulated that maintaining the global production of pre-rRNA avoids activation of nucleolar stress, which can lead to cell cycle arrest and apoptosis [85]. In addition, nucleolar proteins may acquire unique functions to promote viral infection as exemplified for Nucleolin, which is recruited to viral replication compartments (RCs) during HSV-1 and human cytomegalovirus (hCMV) infection [39, 40, 72]. Finally, nucleolar modifications might be associated with functional disturbances or alterations in rRNA transcription and processing, which could in-turn affect protein translation.

Using different approaches, we show that rRNA synthesis continues during the lytic cycle. This finding is in line with previous reports in HSV-1, describing continuous association of UBF with rDNA along with colocalization with the largest RNA pol I subunit RPA43 [43], and ongoing pre-rRNA synthesis at a rate only slightly below that in uninfected cells [47]. However, unlike HSV-1 infection [47], we did not document an altered rRNA maturation pathway during the lytic cycle. In view of the ongoing rRNA transcription, alternative modifications that may affect translation efficiency, accuracy and termination, but may also alter the subsets of mRNAs targeted for translation, may provide an additional layer of infection control.

The changes that take place on rRNA during reactivation are intriguing. Evidence has recently accumulated indicating that the level of snoRNAs and the modifications they guide are altered during the neoplastic process. For instance, a change in a single snoRNA H/ACA, *SNORA24* which guides two pseudouridine, mediates tumor suppression, and ribosomes lacking *SNORA24*-guided Ψ modifications have alterations in their biophysical properties. These findings highlight a role for specific snoRNAs in safeguarding the organism against oncogenic insults, and demonstrate a functional link between H/ACA snoRNAs regulated by RAS and the biophysical properties of ribosomes in cancer [87]. The altered residues observed in our study are mostly located in the functional domains H69 and PTC. Indeed, in yeast, depletion of 1-5 snoRNAs guiding Ψ on PTC were shown to affect folding of the rRNA and translation [88]. Moreover, depletion of combinations of ψ on the yeast H69 located in the inter subunit bridge, which interacts with both A and P site RNAs, affected translation efficiency and fidelity and the level of modification, and was shown to change the translation specificity of the ribosomes [89]. Indeed, recent finding suggest that alterations in ribosome composition, either by changes the ribosomal proteins or the rRNA modifications, could very well represent examples of epitranscriptomic regulation of mRNA translation [78-81, 90, 91]. Thus the rRNA pattern observed under reactivation may preferentially drive the translation of selected viral mRNAs.

In summary, we describe major changes in the nuclear and nucleolar organization that take place during lytic reactivation of KSHV. These changes may signify cellular nucleolar stress responses, but may also occur due to the recruitment of cellular proteins to replication compartments or association with viral proteins that may have major implications on translation of viral and host cell proteins during lytic infection.

## Materials and methods

### Cell culture and transfection

Human renal cell carcinoma SLK and iSLK cells (kindly provided by Don Ganem, Howard Hughes Medical Institute, UCSF, San Francisco, CA, and Rolf Renne, University of Florida, Gainesville, FL) [60] were grown in Dulbecco’s modified Eagle’s medium (DMEM) (Biological Industries, Israel) containing 50 IU/ml penicillin and 50 µg/ ml streptomycin (Biological industries, Israel), and supplemented with 10% heat-inactivated fetal calf serum (FCS) (Biological Industries, Israel). iSLK cells were grown in the presence of 250 µg/ml G418 and 1 µg/ml Puromycin (A.G. Scientific Inc.) to maintain the Tet-on transactivator and the RTA expression cassette, respectively. The growth medium of BAC16-infected iSLK cells was supplemented with 600 µg/ml hygromycin (MegaPharm, San Diego, CA). KSHV-infected iSLK cells were treated with 1 μg/ml Dox and 1 mM sodium butyrate (Sigma), in the absence of hygromycin, puromycin, and G418, to induce RTA transgene expression and lytic cycle reactivation. 0.5 μM Phosphonoacetic acid (PAA) (Sigma) was added to inhibit viral DNA replication.

### Fluorescence and immunofluorescence assay (IFA)

Cells were seeded on coverslips in a 24-well plate, washed with PBS (Biological Industries, Israel), and fixed with 4% formaldehyde in PBS for 25 min at room temperature. After fixation, cells were washed with PBS, and permeabilized and blocked in PBS containing 0.2% Triton X-100% and 1% BSA. Slides were incubated with primary antibodies to UBF (Sigma/Santa Cruz), Fibrillarin (Abcam), NPM (Abcam), Nucleolin (Abcam), RPA194 (Santa Cruz), PICT-1 (Santa Cruz), PML (Santa Cruz), LANA-1 (LN53, Santa Cruz) or ORF59 (a kind gift from Prof. Bala Chandran, University of South Florida) [69] at 40C, followed by incubation with secondary conjugated antibody (Rhodamine, Cy3 or Alexa 647, Jackson ImmunoResearch Laboratories, Inc., West Grove, PA) for 1 hr at room temperature. To stain the nuclei, cells were incubated for 10-min with 0.05 µg/ml Hoechst dye (Sigma) in PBS. The slides were mounted with anti-fading medium (1% n-Propyl gallate, 90% Glycerol in PBS). Cells were examined and photographed under a confocal laser-scanning microscope (Leica SP8 Confocal Live Microscope).

### IFA combined with fluorescence in situ hybridization (FISH)

Cells were washed with PBS, mounted on poly-L-lysine-coated slides, and fixed with 1.6% formaldehyde in PBS at room temperature for 30 min. Cells were then treated with 1% Triton X-100 for 3 min, washed with PBS and prehybridized with hybridization buffer (60% deionized formamide, 50 mM sodium phosphate, pH 7.2, 0.5 mg/ml salmon sperm DNA, 1 µg/ml tRNA, and 5X Denhardt’s solution in 2X SSC) overnight at 720C. ITS-1 RNA probe was prepared by PCR using DNA from HeLa cells as template and forward and reverse primers 5’-CCTGTGGGGTGGTGTC-3’ and 5’-TTAATACGACTCACTATAGGGGTTGCCTCAGGCCG-3’, respectively. The resulting product containing a T7 promoter was cloned into pGEM-T Easy vector and was used to prepare anti-sense RNA probe labelled with biotin-UTP. The probes were diluted in 50% formamide, denatured for 5 min at 850C, and immediately chilled on ice. Next, probes were diluted in hybridization buffer, denatured again for 5 min at 850C, and immediately added to the slides at a final concentration of 2 ng per slide. The slides were covered, sealed, heated for 15 min at 850C, and transferred to 550C for overnight incubation. Slides were washed with 2X SSC for 5 min at room temperature, 0.2X SSC for 30 min at 60°C, and finally with 0.2X SSC for 5 min at room temperature, and then incubated with Alexa Fluor 647 streptavidin conjugate (Molecular Probes). Slides were washed with 0.2X SSC and then incubated with 0.2X SSC containing 10% Bovine Serum Albumin at room temperature for 30 min. Then, cells were incubated antibodies to Fibrillarin for 2-h, washed with 0.2X SSC, and incubated for 1 hr with secondary conjugated antibody (Rhodamine). Nuclei were stained with DAPI, and cells were visualized under Leica SP8 Confocal Live Microscope.

### Reverse-transcription (RT)-quantitative PCR (RT-qPCR)

Total RNA was extracted by using EZ-RNA total RNA isolation kit (Biological Industries, Israel). Residual DNA contamination was eliminated by subsequent treatment with DNase (Turbo DNA-free kit, Ambion). cDNA was generated using 1 µg total RNA with a qScript cDNA synthesis kit (Quanta Biosciences) primed with random hexamers, according to the manufacturer’s instructions. RT-qPCR was performed in a total volume of 10 µl with 1.5 µl of cDNA diluted 1:2, and 0.15 M of each primer specific for rRNA, and Fast Sybr green master mix (Applied Biosystems). The expression levels obtained for each gene were normalized to those of 18S RNA. The primers used were as follows: for *47S* (rRNA), 5’-TGTCAGGCGTTCTCGTCTC-3’ and 5’-GAGAGCACGACGTCACCAC-3’; for *18S*, 5’-ACCGATTGGATGGTTTAGTGAG-3’ and 5’-CCTACGGAAACCTTGTTACGAC-3’. All PCRs were run in triplicate on a StepOne Plus real-time PCR system (Applied Biosystems Inc., Carlsbad, CA).

### Northern blot hybridization

Total RNA was prepared with EZ-RNA total RNA isolation kit (Biological Industries, Israel), and 20 µg samples were loaded and fractionated on a 1.2% agarose, 2.2 M formaldehyde gel. The RNA was visualized with ethidium bromide. Small RNAs were analyzed on a 10% polyacrylamide gel containing 7 M urea. The RNA was transferred to a nylon membrane (Hybond; Amersham Biosciences) and probed with the following γ-32P-end-labeled oligonucleotides: 5.8S rRNA: 5’-TCAGACAGGCGTAGCCCCGGGAGGAACCCG-3’, 18S rRNA: 5’-ATCGGCCCGAGGTTATCTAGAGTCACCAAA-3’, and 28S rRNA: 5’-CCTCTTCGGGGGACGCGCGCGTGGCCCCGA-3’. For labeling, 50 pmoles of each probe was incubated with 50 pmoles of [γ32P]ATP and T4 polynucleotide kinase (Promega) for 30 min at 37°C. In all cases, membranes were washed twice in 2X SSC, 0.1% SDS at 60°C for 20 min, and once in 0.1X SSC, 0.1% SDS at 43°C for 15 min. 18S and 28S rRNA visualized by ethidium bromide was used as loading control.

### BrU labeling

Cells were seeded on coverslips, washed three times with cold PBS (Biological Industries, Israel) and three times with Transcription Buffer (20 mM K-Glutamate, 3 mM MgCl2, 20 mM HEPES pH 7.5, 120 mM Sucrose, 10 mg/ml Leupeptin, 1 mM DTT), and then incubated with Transcription Buffer for 5 min on ice, and permeabilzed with 0.03% Lysolecithin for another 5 minutes. After three washes with Transcription Buffer, the cells were incubated for 30 min at 370C with labeling buffer (0.05 M Creatine phosphate, 4 mM ATP, 2 mM GTP, 2mM CTP, 0.5 mM Br-UTP, CPK, 100 mM DTT, 1 mg/ml Leupeptin in Transcription buffer). Next, 100 µg/ml alpha-amanitin was added to the buffer to block RNA polymerase II transcription. The cells were washed three times with Transcription Buffer and then fixed with 4% formaldehyde in PBS for 25 min at room temperature. After fixation, cells were washed with PBS and stained with BrU antibody and secondary anti-mouse Rhodamin-conjugated antibody in PBS containing 0.2% Triton X-100%. After washing, the cells were blocked with PBS containing 10% Bovine Serum Albumin at room temperature for 30 min and then stained with rabbit anti-UBF for 2h and Cy5-conjugated anti-rabbit secondary antibody for 1-hr. Hoechst staining was used to detect chromatin.

### Mapping the Pseudouridine nucleotides by the CMC method

Pseudouridine sites were mapped as previously described [90]. RNA from BAC16-mCherry-ORF45-infected iSLK cells and SLK cells that were treated with Dox and n-Butyrate for 48-hr was extracted and treated with CMC (N-cyclohexyl-N′-β-(4-methylmorpholinium) ethylcarbodiimide p-tosylate) in CMC buffer (0.17 M CMC in 50 mM bicine, pH 8.3, 4 mM EDTA, 7 M urea) at 370C for 20 min. Under these conditions, the incorporation of CMC in the place of Ψ terminates reverse transcription one nucleotide 3’to the pseudouridylated base, and thereby allows analysis of RNA pseudouridylation at single nucleotide resolution. To remove all the CMC groups except those linked to the Ψ, the CMC-treated RNA was subjected to alkali hydrolysis with Na2CO3 (50 mM, pH 10.4) at 37 °C for 4h. The RNA was then fragmented to a size range of 50-150 nt, adaptor was ligated to the 3′ ends, and cDNA was prepared using reverse transcriptase. Then, an adaptor was ligated to the cDNA, and after amplification, the samples were sequenced in Illumina machine in paired end mode. The reads were mapped to human rRNA using Smalt v0.7.5 (default parameters). Each read pair was “virtually” extended to cover the area from the beginning of the first read to the end of its mate. For each base, the number of reads initializing at that location as well as the number of reads covering the position were calculated. A combination of Bedtools and in-house Perl scripts were used to calculate the Ψ-ratio and Ψ-fc (fold change). For each nucleotide, we computed the Ψ-ratio, dividing the number of reads covering that nucleotide by the number of nucleotides initiating at the following base (i.e. corresponding to the last position copied by the reverse transcriptase). This was repeated for (−CMC) and (+CMC) samples. The Ψ-fc was computed as the log2-fold change of the Ψ-ratios in iSLK versus the SLK samples. We applied this threshold to each sample. Primer extension was performed as previously described [92] with or without CMC and 5′-end-labeled oligonucleotides specific to target rRNAs: Ψ 4636-4628: 5’-CCCACAGATGGTAGCTTCGC-3’, Ψ 4689: 5’-GCCGTATCGTTCCGCCTGGG-3’ ,Ψ 4361: 5’-CAAGCCAGTTATCCCTGTGG-3’. The extension products were analyzed on 12% polyacrylamide– 7 M urea gel, next to sequencing reactions performed using the same primer. Band intensity was quantified using ImageJ software (http://imagej.nih.gov/ij/).

## Acknowledgements

We gratefully acknowledge Prof. Don Ganem and Prof. Bala Chandran for gifts of reagents. This work was supported by a grant from the Israel Science Foundation (Grant no. 1365/15).

## Supplemental Figures and Tables

**<u>Figure S1.</u>** Localization of hyper/hypo-modified Ψ positions on the secondary structure of human rRNA based on four independent replicates of Ψ-seq (Table S1), highlighting different domains of the rRNA: LSU rRNA (A); SSU rRNA (B). The secondary structure of human rRNA was derived from (http://www.rna.ccbb.utexas.edu/).

**<u>Table S1</u>**. **The fold-change of Ψs in (iSLK/SLK) based on Ψ-seq for rRNA**. Ψ-fc(log2) across four independent biological replicates for pseudouridylated sites was calculated for all Ψs. Ψ-fc(log2) >1.3 (iSLK/SLK) in at least two independent replicates was considered hypermodified Ψ. Hypermodified and hypomodified Ψ sites are indicated in red and blue, respectively. snoRNAs predicted to guide the corresponding Ψ positions are indicated on the right column.

## Supplemental Movie Legends

**<u>Movies S1 & S2</u>**. **UBF colocalizes with viral replication compartments during lytic reactivation**. 3D stacks of BAC16-infected iSLK cells that were treated with Dox and n-Butyrate for 48-hr to induce lytic reactivation. Cells were stained with anti-ORF59 and secondary Rhodamine-conjugated antibody, and subsequently with anti-UBF and anti-Rabbit Cy5-cojugated secondary antibody (Cyan). Chromatin was detected by Hoechst staining. Each Z-stack contains 31 planes at 0.5 µm steps.

**<u>Movie S3</u>**. **The distribution of UBF and Fibrillarin during lytic induction**. 3D stacks of BAC16-infected iSLK cells that were treated with Dox and n-Butyrate for 48-hr to induce lytic reactivation. Cells were stained with anti-UBF (Cyan), anti-Fibrillarin (Red) and Hoechst (Blue). Each z-stack contains 31 planes at 0.5 µm steps. The distribution of UBF indicates that lytic virus reactivation took place in both cells.

**<u>Movie S4</u>**. **The distribution of UBF and Nucleolin during lytic induction**. 3D stacks of BAC16-infected iSLK cells that were treated with Dox and n-Butyrate for 48-hr to induce lytic reactivation. Cells were stained with anti-UBF (Red), anti-Nucleolin (Cyan), and Hoechst (Blue). Each z-stack contains 31 planes at 0.5 µm steps. The distribution of UBF indicates that lytic virus reactivation took place in the upper cell only.

## References

1. Chang Y, Cesarman E, Pessin MS, Lee F, Culpepper J, Knowles DM, et al. Identification of herpesvirus-like DNA sequences in AIDS-associated Kaposi’s sarcoma. Science. 1994;266(5192):1865–9.

2. Mesri EA, Cesarman E, Boshoff C. Kaposi’s sarcoma and its associated herpesvirus. Nat Rev Cancer. 2010;10(10):707–19. doi: 10.1038/nrc2888. PubMed PMID: 20865011.

3. Dittmer DP, Damania B. Kaposi sarcoma associated herpesvirus pathogenesis (KSHV)--an update. Curr Opin Virol. 2013;3(3):238–44. doi: 10.1016/j.coviro.2013.05.012. PubMed PMID: 23769237; PubMed Central PMCID: PMC3716290.

4. Gramolelli S, Schulz TF. The role of Kaposi sarcoma-associated herpesvirus in the pathogenesis of Kaposi sarcoma. J Pathol. 2015;235(2):368-80. Epub 2014/09/13. doi: 10.1002/path.4441. PubMed PMID: 25212381.

5. Goncalves PH, Ziegelbauer J, Uldrick TS, Yarchoan R. Kaposi sarcoma herpesvirus-associated cancers and related diseases. Curr Opin HIV AIDS. 2017;12(1):47–56. doi: 10.1097/COH.0000000000000330. PubMed PMID: 27662501.

6. Yan L, Majerciak V, Zheng ZM, Lan K. Towards Better Understanding of KSHV Life Cycle: from Transcription and Posttranscriptional Regulations to Pathogenesis. Virol Sin. 2019;34(2):135-61. Epub 2019/04/27. doi: 10.1007/s12250-019-00114-3. PubMed PMID: 31025296; PubMed Central PMCID: PMCPMC6513836.

7. Carroll PA, Brazeau E, Lagunoff M. Kaposi’s sarcoma-associated herpesvirus infection of blood endothelial cells induces lymphatic differentiation. Virology. 2004;328(1):7–18.

8. Chen CP, Lyu Y, Chuang F, Nakano K, Izumiya C, Jin D, et al. Kaposi’s Sarcoma-Associated Herpesvirus Hijacks RNA Polymerase II To Create a Viral Transcriptional Factory. J Virol. 2017;91(11). Epub 2017/03/24. doi: 10.1128/JVI.02491-16. PubMed PMID: 28331082; PubMed Central PMCID: PMCPMC5432858.

9. Vallery TK, Withers JB, Andoh JA, Steitz JA. Kaposi’s Sarcoma-Associated Herpesvirus mRNA Accumulation in Nuclear Foci Is Influenced by Viral DNA Replication and Viral Noncoding Polyadenylated Nuclear RNA. J Virol. 2018;92(13). Epub 2018/04/13. doi: 10.1128/JVI.00220-18. PubMed PMID: 29643239; PubMed Central PMCID: PMCPMC6002709.

10. Taylor TJ, McNamee EE, Day C, Knipe DM. Herpes simplex virus replication compartments can form by coalescence of smaller compartments. Virology. 2003;309(2):232-47. Epub 2003/05/22. doi: 10.1016/s0042-6822(03)00107-7. PubMed PMID: 12758171.

11. Monier K, Armas JC, Etteldorf S, Ghazal P, Sullivan KF. Annexation of the interchromosomal space during viral infection. Nat Cell Biol. 2000;2(9):661-5. Epub 2000/09/12. doi: 10.1038/35023615. PubMed PMID: 10980708.

12. Simpson-Holley M, Colgrove RC, Nalepa G, Harper JW, Knipe DM. Identification and functional evaluation of cellular and viral factors involved in the alteration of nuclear architecture during herpes simplex virus 1 infection. J Virol. 2005;79(20):12840-51. Epub 2005/09/29. doi: 10.1128/JVI.79.20.12840-12851.2005. PubMed PMID: 16188986; PubMed Central PMCID: PMCPMC1235858.

13. Zakaryan H, Stamminger T. Nuclear remodelling during viral infections. Cell Microbiol. 2011;13(6):806-13. Epub 2011/04/20. doi: 10.1111/j.1462-5822.2011.01596.x. PubMed PMID: 21501365.

14. Boisvert FM, van Koningsbruggen S, Navascues J, Lamond AI. The multifunctional nucleolus. NatRevMolCell Biol. 2007;8(7):574–85.

15. Pederson T. The nucleolus. Cold Spring Harb Perspect Biol. 2011;3(3). Epub 2010/11/26. doi: 10.1101/cshperspect.a000638. PubMed PMID: 21106648; PubMed Central PMCID: PMCPMC3039934.

16. Nemeth A, Grummt I. Dynamic regulation of nucleolar architecture. Curr Opin Cell Biol. 2018;52:105-11. Epub 2018/03/13. doi: 10.1016/j.ceb.2018.02.013. PubMed PMID: 29529563.

17. Iarovaia OV, Minina EP, Sheval EV, Onichtchouk D, Dokudovskaya S, Razin SV, et al. Nucleolus: A Central Hub for Nuclear Functions. Trends Cell Biol. 2019;29(8):647-59. Epub 2019/06/10. doi: 10.1016/j.tcb.2019.04.003. PubMed PMID: 31176528.

18. Correll CC, Bartek J, Dundr M. The Nucleolus: A Multiphase Condensate Balancing Ribosome Synthesis and Translational Capacity in Health, Aging and Ribosomopathies. Cells. 2019;8(8). Epub 2019/08/14. doi: 10.3390/cells8080869. PubMed PMID: 31405125; PubMed Central PMCID: PMCPMC6721831.

19. Tsekrekou M, Stratigi K, Chatzinikolaou G. The Nucleolus: In Genome Maintenance and Repair. Int J Mol Sci. 2017;18(7). Epub 2017/07/04. doi: 10.3390/ijms18071411. PubMed PMID: 28671574; PubMed Central PMCID: PMCPMC5535903.

20. Stepinski D. The nucleolus, an ally, and an enemy of cancer cells. Histochem Cell Biol. 2018;150(6):607-29. Epub 2018/08/15. doi: 10.1007/s00418-018-1706-5. PubMed PMID: 30105457; PubMed Central PMCID: PMCPMC6267422.

21. Hiscox JA. The nucleolus--a gateway to viral infection? Arch Virol. 2002;147(6):1077-89. Epub 2002/07/12. doi: 10.1007/s00705-001-0792-0. PubMed PMID: 12111420.

22. Greco A. Involvement of the nucleolus in replication of human viruses. Rev Med Virol. 2009;19(4):201-14. Epub 2009/04/29. doi: 10.1002/rmv.614. PubMed PMID: 19399920.

23. Hiscox JA, Whitehouse A, Matthews DA. Nucleolar proteomics and viral infection. Proteomics. 2010;10(22):4077-86. Epub 2010/07/28. doi: 10.1002/pmic.201000251. PubMed PMID: 20661956.

24. Salvetti A, Greco A. Viruses and the nucleolus: the fatal attraction. Biochim Biophys Acta. 2014;1842(6):840–7. doi: 10.1016/j.bbadis.2013.12.010. PubMed PMID: 24378568.

25. Glingston RS, Deb R, Kumar S, Nagotu S. Organelle dynamics and viral infections: at cross roads. Microbes Infect. 2019;21(1):20-32. Epub 2018/06/29. doi: 10.1016/j.micinf.2018.06.002. PubMed PMID: 29953921.

26. Rawlinson SM, Moseley GW. The nucleolar interface of RNA viruses. Cell Microbiol. 2015;17(8):1108-20. Epub 2015/06/05. doi: 10.1111/cmi.12465. PubMed PMID: 26041433.

27. Pyper JM, Clements JE, Zink MC. The nucleolus is the site of Borna disease virus RNA transcription and replication. J Virol. 1998;72(9):7697-702. Epub 1998/08/08. PubMed PMID: 9696879; PubMed Central PMCID: PMCPMC110048.

28. Sonntag F, Schmidt K, Kleinschmidt JA. A viral assembly factor promotes AAV2 capsid formation in the nucleolus. Proc Natl Acad Sci U S A. 2010;107(22):10220-5. Epub 2010/05/19. doi: 10.1073/pnas.1001673107. PubMed PMID: 20479244; PubMed Central PMCID: PMCPMC2890453.

29. Kubota S, Siomi H, Satoh T, Endo S, Maki M, Hatanaka M. Functional similarity of HIV-I rev and HTLV-I rex proteins: identification of a new nucleolar-targeting signal in rev protein. Biochem Biophys Res Commun. 1989;162(3):963-70. Epub 1989/08/15. doi: 10.1016/0006-291x(89)90767-5. PubMed PMID: 2788417.

30. Cochrane AW, Perkins A, Rosen CA. Identification of sequences important in the nucleolar localization of human immunodeficiency virus Rev: relevance of nucleolar localization to function. J Virol. 1990;64(2):881-5. Epub 1990/02/01. PubMed PMID: 2404140; PubMed Central PMCID: PMCPMC249184.

31. Waggoner S, Sarnow P. Viral ribonucleoprotein complex formation and nucleolar-cytoplasmic relocalization of nucleolin in poliovirus-infected cells. J Virol. 1998;72(8):6699-709. Epub 1998/07/11. PubMed PMID: 9658117; PubMed Central PMCID: PMCPMC109870.

32. Banerjee R, Weidman MK, Navarro S, Comai L, Dasgupta A. Modifications of both selectivity factor and upstream binding factor contribute to poliovirus-mediated inhibition of RNA polymerase I transcription. J Gen Virol. 2005;86(Pt 8):2315-22. Epub 2005/07/22. doi: 10.1099/vir.0.80817-0. PubMed PMID: 16033979.

33. Gouzil J, Fablet A, Lara E, Caignard G, Cochet M, Kundlacz C, et al. Nonstructural Protein NSs of Schmallenberg Virus Is Targeted to the Nucleolus and Induces Nucleolar Disorganization. J Virol. 2017;91(1). Epub 2016/11/01. doi: 10.1128/JVI.01263-16. PubMed PMID: 27795408; PubMed Central PMCID: PMCPMC5165206.

34. Matthews DA. Adenovirus protein V induces redistribution of nucleolin and B23 from nucleolus to cytoplasm. J Virol. 2001;75(2):1031-8. Epub 2001/01/03. doi: 10.1128/JVI.75.2.1031-1038.2001. PubMed PMID: 11134316; PubMed Central PMCID: PMCPMC113999.

35. Okuwaki M, Iwamatsu A, Tsujimoto M, Nagata K. Identification of nucleophosmin/B23, an acidic nucleolar protein, as a stimulatory factor for in vitro replication of adenovirus DNA complexed with viral basic core proteins. J Mol Biol. 2001;311(1):41-55. Epub 2001/07/27. doi: 10.1006/jmbi.2001.4812. PubMed PMID: 11469856.

36. Samad MA, Okuwaki M, Haruki H, Nagata K. Physical and functional interaction between a nucleolar protein nucleophosmin/B23 and adenovirus basic core proteins. FEBS Lett. 2007;581(17):3283-8. Epub 2007/07/03. doi: 10.1016/j.febslet.2007.06.024. PubMed PMID: 17602943.

37. Ugai H, Dobbins GC, Wang M, Le LP, Matthews DA, Curiel DT. Adenoviral protein V promotes a process of viral assembly through nucleophosmin 1. Virology. 2012;432(2):283-95. Epub 2012/06/22. doi: 10.1016/j.virol.2012.05.028. PubMed PMID: 22717133; PubMed Central PMCID: PMCPMC3423539.

38. Lawrence FJ, McStay B, Matthews DA. Nucleolar protein upstream binding factor is sequestered into adenovirus DNA replication centres during infection without affecting RNA polymerase I location or ablating rRNA synthesis. J Cell Sci. 2006;119(Pt 12):2621-31. Epub 2006/06/10. doi: 10.1242/jcs.02982. PubMed PMID: 16763197.

39. Genoveso MJ, Hisaoka M, Komatsu T, Wodrich H, Nagata K, Okuwaki M. Formation of adenovirus DNA replication compartments and viral DNA accumulation sites by host chromatin regulatory proteins including NPM1. FEBS J. 2020;287(1):205-17. Epub 2019/08/01. doi: 10.1111/febs.15027. PubMed PMID: 31365788.

40. Calle A, Ugrinova I, Epstein AL, Bouvet P, Diaz JJ, Greco A. Nucleolin is required for an efficient herpes simplex virus type 1 infection. J Virol. 2008;82(10):4762–73. doi: 10.1128/JVI.00077-08. PubMed PMID: 18321972; PubMed Central PMCID: PMC2346767.

41. Stow ND, Evans VC, Matthews DA. Upstream-binding factor is sequestered into herpes simplex virus type 1 replication compartments. J Gen Virol. 2009;90(Pt 1):69-73. Epub 2008/12/18. doi: 10.1099/vir.0.006353-0. PubMed PMID: 19088274; PubMed Central PMCID: PMCPMC2885023.

42. Lymberopoulos MH, Pearson A. Involvement of UL24 in herpes-simplex-virus-1-induced dispersal of nucleolin. Virology. 2007;363(2):397–409. doi: 10.1016/j.virol.2007.01.028. PubMed PMID: 17346762.

43. Lymberopoulos MH, Pearson A. Relocalization of upstream binding factor to viral replication compartments is UL24 independent and follows the onset of herpes simplex virus 1 DNA synthesis. J Virol. 2010;84(9):4810–5. doi: 10.1128/JVI.02437-09. PubMed PMID: 20147409; PubMed Central PMCID: PMC2863781.

44. Lymberopoulos MH, Bourget A, Ben Abdeljelil N, Pearson A. Involvement of the UL24 protein in herpes simplex virus 1-induced dispersal of B23 and in nuclear egress. Virology. 2011;412(2):341–8. doi: 10.1016/j.virol.2011.01.016. PubMed PMID: 21316727.

45. Simonin D, Diaz JJ, Masse T, Madjar JJ. Persistence of ribosomal protein synthesis after infection of HeLa cells by herpes simplex virus type 1. J Gen Virol. 1997;78 (Pt 2):435–43. doi: 10.1099/0022-1317-78-2-435. PubMed PMID: 9018067.

46. Besse S, Puvion-Dutilleul F. Intranuclear retention of ribosomal RNAs in response to herpes simplex virus type 1 infection. J Cell Sci. 1996;109 (Pt 1):119-29. Epub 1996/01/01. PubMed PMID: 8834797.

47. Belin S, Kindbeiter K, Hacot S, Albaret MA, Roca-Martinez JX, Therizols G, et al. Uncoupling ribosome biogenesis regulation from RNA polymerase I activity during herpes simplex virus type 1 infection. RNA. 2010;16(1):131–40. doi: 10.1261/rna.1935610. PubMed PMID: 19934231; PubMed Central PMCID: PMC2802023.

48. Reyes ED, Kulej K, Pancholi NJ, Akhtar LN, Avgousti DC, Kim ET, et al. Identifying Host Factors Associated with DNA Replicated During Virus Infection. Mol Cell Proteomics. 2017;16(12):2079-97. Epub 2017/10/04. doi: 10.1074/mcp.M117.067116. PubMed PMID: 28972080; PubMed Central PMCID: PMCPMC5724173.

49. Sadagopan S, Sharma-Walia N, Veettil MV, Bottero V, Levine R, Vart RJ, et al. Kaposi’s sarcoma-associated herpesvirus upregulates angiogenin during infection of human dermal microvascular endothelial cells, which induces 45S rRNA synthesis, antiapoptosis, cell proliferation, migration, and angiogenesis. J Virol. 2009;83(7):3342-64. Epub 2009/01/23. doi: 10.1128/JVI.02052-08. PubMed PMID: 19158252; PubMed Central PMCID: PMCPMC2655544.

50. Sarek G, Jarviluoma A, Moore HM, Tojkander S, Vartia S, Biberfeld P, et al. Nucleophosmin phosphorylation by v-cyclin-CDK6 controls KSHV latency. PLoS Pathog. 2010;6(3):e1000818. Epub 2010/03/25. doi: 10.1371/journal.ppat.1000818. PubMed PMID: 20333249; PubMed Central PMCID: PMCPMC2841626.

51. Muller M, Hutin S, Marigold O, Li KH, Burlingame A, Glaunsinger BA. A ribonucleoprotein complex protects the interleukin-6 mRNA from degradation by distinct herpesviral endonucleases. PLoS Pathog. 2015;11(5):e1004899. Epub 2015/05/13. doi: 10.1371/journal.ppat.1004899. PubMed PMID: 25965334; PubMed Central PMCID: PMCPMC4428876.

52. Boyne JR, Whitehouse A. Nucleolar trafficking is essential for nuclear export of intronless herpesvirus mRNA. Proc Natl Acad Sci U S A. 2006;103(41):15190-5. Epub 2006/09/29. doi: 10.1073/pnas.0604890103. PubMed PMID: 17005724; PubMed Central PMCID: PMCPMC1622798.

53. Jackson BR, Noerenberg M, Whitehouse A. The Kaposi’s Sarcoma-Associated Herpesvirus ORF57 Protein and Its Multiple Roles in mRNA Biogenesis. Front Microbiol. 2012;3:59. Epub 2012/03/01. doi: 10.3389/fmicb.2012.00059. PubMed PMID: 22363332; PubMed Central PMCID: PMCPMC3282479.

54. Cheng EH, Nicholas J, Bellows DS, Hayward GS, Guo HG, Reitz MS, et al. A Bcl-2 homolog encoded by Kaposi sarcoma-associated virus, human herpesvirus 8, inhibits apoptosis but does not heterodimerize with Bax or Bak. ProcNatlAcadSciUSA. 1997;94(2):690–4.

55. Sarid R, Sato T, Bohenzky RA, Russo JJ, Chang Y. Kaposi’s sarcoma-associated herpesvirus encodes a functional bcl-2 homologue. NatMed. 1997;3(3):293–8.

56. Cuconati A, White E. Viral homologs of BCL-2: role of apoptosis in the regulation of virus infection. Genes Dev. 2002;16(19):2465–78.

57. Kalt I, Borodianskiy-Shteinberg T, Schachor A, Sarid R. GLTSCR2/PICT-1, a putative tumor suppressor gene product, induces the nucleolar targeting of the Kaposi’s sarcoma-associated herpesvirus KS-Bcl-2 protein. J Virol. 2010;84(6):2935–45. doi: 10.1128/JVI.00757-09. PubMed PMID: 20042497; PubMed Central PMCID: PMC2826064.

58. Bussey KA, Lau U, Schumann S, Gallo A, Osbelt L, Stempel M, et al. The interferon-stimulated gene product oligoadenylate synthetase-like protein enhances replication of Kaposi’s sarcoma-associated herpesvirus (KSHV) and interacts with the KSHV ORF20 protein. PLoS Pathog. 2018;14(3):e1006937. Epub 2018/03/03. doi: 10.1371/journal.ppat.1006937. PubMed PMID: 29499066; PubMed Central PMCID: PMCPMC5851652.

59. Wang Y, Li H, Tang Q, Maul GG, Yuan Y. Kaposi’s sarcoma-associated herpesvirus ori-Lyt-dependent DNA replication: involvement of host cellular factors. J Virol. 2008;82(6):2867–82. doi: 10.1128/JVI.01319-07. PubMed PMID: 18199640; PubMed Central PMCID: PMC2259006.

60. Myoung J, Ganem D. Generation of a doxycycline-inducible KSHV producer cell line of endothelial origin: maintenance of tight latency with efficient reactivation upon induction. J Virol Methods. 2011;174(1-2):12–21. doi: 10.1016/j.jviromet.2011.03.012. PubMed PMID: 21419799; PubMed Central PMCID: PMC3095772.

61. Brulois KF, Chang H, Lee AS, Ensser A, Wong LY, Toth Z, et al. Construction and manipulation of a new Kaposi’s sarcoma-associated herpesvirus bacterial artificial chromosome clone. J Virol. 2012;86(18):9708–20. doi: 10.1128/JVI.01019-12. PubMed PMID: 22740391; PubMed Central PMCID: PMC3446615.

62. Bergson S, Kalt I, Itzhak I, Brulois KF, Jung JU, Sarid R. Fluorescent tagging and cellular distribution of the Kaposi’s sarcoma-associated herpesvirus ORF45 tegument protein. J Virol. 2014;88(21):12839–52. doi: 10.1128/JVI.01091-14. PubMed PMID: 25165104.

63. Lallemand-Breitenbach V, de The H. PML nuclear bodies: from architecture to function. Curr Opin Cell Biol. 2018;52:154-61. Epub 2018/05/04. doi: 10.1016/j.ceb.2018.03.011. PubMed PMID: 29723661.

64. Full F, Jungnickl D, Reuter N, Bogner E, Brulois K, Scholz B, et al. Kaposi’s sarcoma associated herpesvirus tegument protein ORF75 is essential for viral lytic replication and plays a critical role in the antagonization of ND10-instituted intrinsic immunity. PLoS Pathog. 2014;10(1):e1003863. Epub 2014/01/24. doi: 10.1371/journal.ppat.1003863. PubMed PMID: 24453968; PubMed Central PMCID: PMCPMC3894210.

65. Full F, Ensser A. Early Nuclear Events after Herpesviral Infection. J Clin Med. 2019;8(9). Epub 2019/09/11. doi: 10.3390/jcm8091408. PubMed PMID: 31500286; PubMed Central PMCID: PMCPMC6780142.

66. Weidner-Glunde M, Mariggio G, Schulz TF. Kaposi’s Sarcoma-Associated Herpesvirus Latency-Associated Nuclear Antigen: Replicating and Shielding Viral DNA during Viral Persistence. J Virol. 2017;91(14). Epub 2017/04/28. doi: 10.1128/JVI.01083-16. PubMed PMID: 28446671; PubMed Central PMCID: PMCPMC5487577.

67. De Leo A, Calderon A, Lieberman PM. Control of Viral Latency by Episome Maintenance Proteins. Trends Microbiol. 2019. Epub 2019/10/19. doi: 10.1016/j.tim.2019.09.002. PubMed PMID: 31624007.

68. Bertrand L, Pearson A. The conserved N-terminal domain of herpes simplex virus 1 UL24 protein is sufficient to induce the spatial redistribution of nucleolin. J Gen Virol. 2008;89(Pt 5):1142-51. Epub 2008/04/19. doi: 10.1099/vir.0.83573-0. PubMed PMID: 18420791.

69. Chan SR, Bloomer C, Chandran B. Identification and characterization of human herpesvirus-8 lytic cycle-associated ORF 59 protein and the encoding cDNA by monoclonal antibody. Virology. 1998;240(1):118–26.

70. Chan SR, Chandran B. Characterization of human herpesvirus 8 ORF59 protein (PF-8) and mapping of the processivity and viral DNA polymerase-interacting domains. J Virol. 2000;74(23):10920-9. Epub 2000/11/09. doi: 10.1128/jvi.74.23.10920-10929.2000. PubMed PMID: 11069986; PubMed Central PMCID: PMCPMC113171.

71. Zhou X, Liao Q, Ricciardi RP, Peng C, Chen X. Kaposi’s sarcoma-associated herpesvirus processivity factor-8 dimerizes in cytoplasm before being translocated to nucleus. Biochem Biophys Res Commun. 2010;397(3):520-5. Epub 2010/06/03. doi: 10.1016/j.bbrc.2010.05.147. PubMed PMID: 20515658.

72. Strang BL, Boulant S, Coen DM. Nucleolin associates with the human cytomegalovirus DNA polymerase accessory subunit UL44 and is necessary for efficient viral replication. J Virol. 2010;84(4):1771-84. Epub 2009/12/17. doi: 10.1128/JVI.01510-09. PubMed PMID: 20007282; PubMed Central PMCID: PMCPMC2812382.

73. Strang BL, Boulant S, Kirchhausen T, Coen DM. Host cell nucleolin is required to maintain the architecture of human cytomegalovirus replication compartments. MBio. 2012;3(1). Epub 2012/02/10. doi: 10.1128/mBio.00301-11. PubMed PMID: 22318319; PubMed Central PMCID: PMCPMC3280463.

74. Sloan KE, Bohnsack MT, Watkins NJ. The 5S RNP couples p53 homeostasis to ribosome biogenesis and nucleolar stress. Cell Rep. 2013;5(1):237-47. Epub 2013/10/15. doi: 10.1016/j.celrep.2013.08.049. PubMed PMID: 24120868; PubMed Central PMCID: PMCPMC3808153.

75. Cepeda LPP, Bagatelli FFM, Santos RM, Santos MDM, Nogueira FCS, Oliveira CC. The ribosome assembly factor Nop53 controls association of the RNA exosome with pre-60S particles inyeast. J Biol Chem. 2019. Epub 2019/10/31. doi: 10.1074/jbc.RA119.010193. PubMed PMID: 31662437.

76. Sloan KE, Warda AS, Sharma S, Entian KD, Lafontaine DLJ, Bohnsack MT. Tuning the ribosome: The influence of rRNA modification on eukaryotic ribosome biogenesis and function. RNA Biol. 2017;14(9):1138-52. Epub 2016/12/03. doi: 10.1080/15476286.2016.1259781. PubMed PMID: 27911188; PubMed Central PMCID: PMCPMC5699541.

77. Decatur WA, Fournier MJ. rRNA modifications and ribosome function. Trends Biochem Sci. 2002;27(7):344-51. Epub 2002/07/13. doi: 10.1016/s0968-0004(02)02109-6. PubMed PMID: 12114023.

78. Dimitrova DG, Teysset L, Carre C. RNA 2’-O-Methylation (Nm) Modification in Human Diseases. Genes (Basel). 2019;10(2). Epub 2019/02/16. doi: 10.3390/genes10020117. PubMed PMID: 30764532; PubMed Central PMCID: PMCPMC6409641.

79. Monaco PL, Marcel V, Diaz JJ, Catez F. 2’-O-Methylation of Ribosomal RNA: Towards an Epitranscriptomic Control of Translation? Biomolecules. 2018;8(4). Epub 2018/10/05. doi: 10.3390/biom8040106. PubMed PMID: 30282949; PubMed Central PMCID: PMCPMC6316387.

80. Ayadi L, Galvanin A, Pichot F, Marchand V, Motorin Y. RNA ribose methylation (2’-O-methylation): Occurrence, biosynthesis and biological functions. Biochim Biophys Acta Gene Regul Mech. 2019;1862(3):253-69. Epub 2018/12/21. doi: 10.1016/j.bbagrm.2018.11.009. PubMed PMID: 30572123.

81. Penzo M, Montanaro L. Turning Uridines around: Role of rRNA Pseudouridylation in Ribosome Biogenesis and Ribosomal Function. Biomolecules. 2018;8(2). Epub 2018/06/08. doi: 10.3390/biom8020038. PubMed PMID: 29874862; PubMed Central PMCID: PMCPMC6023024.

82. Bakin A, Ofengand J. Four newly located pseudouridylate residues in Escherichia coli 23S ribosomal RNA are all at the peptidyltransferase center: analysis by the application of a new sequencing technique. Biochemistry. 1993;32(37):9754-62. Epub 1993/09/21. doi: 10.1021/bi00088a030. PubMed PMID: 8373778.

83. Schwartz S, Bernstein DA, Mumbach MR, Jovanovic M, Herbst RH, Leon-Ricardo BX, et al. Transcriptome-wide mapping reveals widespread dynamic-regulated pseudouridylation of ncRNA and mRNA. Cell. 2014;159(1):148-62. Epub 2014/09/16. doi: 10.1016/j.cell.2014.08.028. PubMed PMID: 25219674; PubMed Central PMCID: PMCPMC4180118.

84. Tusup M, Kundig T, Pascolo S. Epitranscriptomics of cancer. World J Clin Oncol. 2018;9(3):42-55. Epub 2018/06/15. doi: 10.5306/wjco.v9.i3.42. PubMed PMID: 29900123; PubMed Central PMCID: PMCPMC5997933.

85. Boulon S, Westman BJ, Hutten S, Boisvert FM, Lamond AI. The nucleolus under stress. Mol Cell. 2010;40(2):216–27. doi: 10.1016/j.molcel.2010.09.024. PubMed PMID: 20965417; PubMed Central PMCID: PMC2987465.

86. Meng W, Han SC, Li CC, Dong HJ, Chang JY, Wang HR, et al. Cytoplasmic Translocation of Nucleolar Protein NOP53 Promotes Viral Replication by Suppressing Host Defense. Viruses. 2018;10(4). Epub 2018/04/21. doi: 10.3390/v10040208. PubMed PMID: 29677136; PubMed Central PMCID: PMCPMC5923502.

87. McMahon M, Contreras A, Holm M, Uechi T, Forester CM, Pang X, et al. A single H/ACA small nucleolar RNA mediates tumor suppression downstream of oncogenic RAS. Elife. 2019;8. Epub 2019/09/04. doi: 10.7554/eLife.48847. PubMed PMID: 31478838; PubMed Central PMCID: PMCPMC6776443.

88. King TH, Liu B, McCully RR, Fournier MJ. Ribosome structure and activity are altered in cells lacking snoRNPs that form pseudouridines in the peptidyl transferase center. Mol Cell. 2003;11(2):425-35. Epub 2003/03/07. doi: 10.1016/s1097-2765(03)00040-6. PubMed PMID: 12620230.

89. Liang XH, Liu Q, Fournier MJ. rRNA modifications in an intersubunit bridge of the ribosome strongly affect both ribosome biogenesis and activity. Mol Cell. 2007;28(6):965-77. Epub 2007/12/27. doi: 10.1016/j.molcel.2007.10.012. PubMed PMID: 18158895.

90. Chikne V, Doniger T, Rajan KS, Bartok O, Eliaz D, Cohen-Chalamish S, et al. A pseudouridylation switch in rRNA is implicated in ribosome function during the life cycle of Trypanosoma brucei. Sci Rep. 2016;6:25296. Epub 2016/05/05. doi: 10.1038/srep25296. PubMed PMID: 27142987; PubMed Central PMCID: PMCPMC4855143.

91. Genuth NR, Barna M. The Discovery of Ribosome Heterogeneity and Its Implications for Gene Regulation and Organismal Life. Mol Cell. 2018;71(3):364-74. Epub 2018/08/04. doi: 10.1016/j.molcel.2018.07.018. PubMed PMID: 30075139; PubMed Central PMCID: PMCPMC6092941.

92. Mandelboim M, Barth S, Biton M, Liang XH, Michaeli S. Silencing of Sm proteins in Trypanosoma brucei by RNA interference captured a novel cytoplasmic intermediate in spliced leader RNA biogenesis. J Biol Chem. 2003;278(51):51469-78. Epub 2003/10/09. doi: 10.1074/jbc.M308997200. PubMed PMID: 14532264.

